# The G2-phase enriched lncRNA *SNHG26* is necessary for proper cell cycle progression and proliferation

**DOI:** 10.1101/2021.02.22.432245

**Authors:** Helle Samdal, Siv A. Hegre, Konika Chawla, Nina-Beate Liabakk, Per A. Aas, Bjørnar Sporsheim, Pål Sætrom

## Abstract

Long noncoding RNAs (lncRNAs) are involved in the regulation of cell cycle, although only a few have been functionally characterized. By combining RNA sequencing and ChIP sequencing of cell cycle synchronized HaCaT cells we have previously identified lncRNAs highly enriched for cell cycle functions. Based on a cyclic expression profile and an overall high correlation to histone 3 lysine 4 trimethylation (H3K4me3) and RNA polymerase II (Pol II) signals, the lncRNA *SNHG26* was identified as a top candidate. In the present study we report that downregulation of *SNHG26* affects mitochondrial stress, proliferation, cell cycle phase distribution, and gene expression in *cis*- and in *trans*, and that this effect is reversed by upregulation of *SNHG26*. We also find that the effect on cell cycle phase distribution is cell type specific and stable over time. Results indicate an oncogenic role of *SNHG26*, possibly by affecting cell cycle progression through the regulation of downstream MYC-responsive genes.

## Introduction

LncRNAs are more than 200 nucleotides long with none or little protein coding potential. Most lncRNAs are transcribed by Pol II, polyadenylated, and spliced (Chen, 2016). LncRNAs can interact with DNA, RNA, and proteins, and are important regulators of gene expression at the epigenetic, transcriptional, and translational level. Among the different modes of mechanisms used by lncRNAs are the mediation of inter-chromosomal interactions, formation of nuclear bodies, and facilitating changes in histone modifications (Bohmdorfer and Wierzbicki, 2015). LncRNAs can further act as sponges to miRNAs, as scaffolds for chromatin modifying complexes, and as a guide or decoy of transcription factors (Marchese et al., 2017). To date, the Encyclopedia of DNA elements (ENCODE) project (GENCODE v35) has annotated 17957 lncRNA genes which give rise to 46977 different transcripts.

Compared to mRNAs, lncRNAs have a lower and more cell type-specific expression, with some being restricted to only a single cell type. There are also ubiquitously expressed lncRNAs that typically are involved in basal cellular functions that are essential for the cell’s existence (Jiang et al., 2016). LncRNAs are involved in several cellular processes through a variety of mechanisms, and an aberrant expression of lncRNAs is associated with the pathogenesis of several diseases, including cancers where they are involved in tumorigenesis and progression. Specifically, several lncRNAs are important in the regulation of the cell cycle where they are involved in controlling the level of cell cycle regulators such as cyclins, cyclin-dependent kinases (CDKs), CDK inhibitors (CKIs), cell division cycle 25 (*CDC25*), E2 factor (*E2F*), and retinoblastoma (*RB1*) (Kitagawa et al., 2013).

One specific class of lncRNAs are small nucleolar RNA (snoRNA) host genes (SNHG). SnoRNAs consist of 60-300 nucleotides, accumulate in the nucleoli, and are involved with ribosomal RNA (rRNA) processing and the regulation of gene expression (Liang et al., 2019). SnoRNAs are generated from introns only, but if the full-length spliced transcript including exons exist as a stable molecule, it is classified as an SNHG. SNHGs have oncogenic properties connected to proliferation, invasion, metastasis, and cell cycle progression, are often overexpressed in cancers, and may represent biomarkers for disease progression and novel therapeutic targets (Zimta et al., 2020, Williams and Farzaneh, 2012). For nuclear localized SNHGs, the main molecular mechanisms previously reported are modulation of methylation enzymes affecting the DNAs methylation state, and repression of gene transcription through the interaction with transcription factors. Meanwhile, SNHGs located in the cytoplasm have functions like miRNA sponging, repression of translation through direct binding of mRNAs, and interference with protein ubiquitination (Zimta et al., 2020).

*SNHG26* is a multi-exonic gene that hosts the snoRNA *SNORD93*, which can be further processed into a small nucleolar RNA-derived RNA (sdRNA-93) with miRNA-like functions associated with malignancy of breast cancer (Patterson et al., 2017). Previous studies have identified *SNHG26* as dysregulated in cancers including multiple myeloma, bladder, and lung cancer (Liu et al., 2019, Bao et al., 2017, Hu et al., 2017).

In a previous study we combined RNA sequencing (RNA-seq) with ChIP sequencing (ChIP-seq) of cell cycle synchronized HaCaT cells and identified 59 lncRNAs with cell cycle-dependent expression and correlated changes in Pol II occupancy or promoter activity as measured by H3K4me3 and histone 3 lysine 27 trimethylation (H3K27me3) signals (preprint: https://doi.org/10.1101/2021.02.12.430890). Of these cell cycle-associated lncRNAs, *SNHG26* (also known as *AC005682.5*) had the highest overall correlation, and its biological function was investigated by siRNA-mediated knockdown followed by cell cycle- and proliferation assays. Downregulation of *SNHG26* resulted in reduced proliferation in HaCaT, A549, LS411N, and DLD1 cells. We also observed a reduction of HaCaT cells present in the G1 phase and an enrichment of cells present in the G2/M phase of the cell cycle in response to *SNHG26* knockdown, indicating a possible role in the G2 to M progression (preprint: https://doi.org/10.1101/2021.02.12.430890).

In the present study, we further characterized the biological function and provide some mechanistic insight into the oncogenic properties of *SNHG26*. We used siRNA, ASO, or CRISPR interference and CRISPR activation (CRISPRi and CRISPRa, respectively) to modulate the expression of *SNHG26* to evaluate its *cis*- and *trans* regulatory abilities and to investigate how it affects proliferation, mitochondrial stress, and cell cycle phase distribution. We used RNA FISH to determine the subcellular localization of *SNHG26* to be mainly nuclear in HaCaT, A549, and LS411N cells, with high-intensity spots surrounded by nuclear and extranuclear low-intensity spots. We report that *SNHG26* affects cell cycle phase distribution in all three investigated cell lines, and that the phase distribution is stable and cell-type specific. Moreover, downregulation of *SNHG26* reduced the viability and caused increased mitochondrial stress. We also find that *SNHG26* has cis-regulatory abilities as it regulates the expression of its neighbor mRNA *TOMM7*, and that this effect is post-transcriptional. Furthermore, CRISPR-mediated upregulation of *SNHG26* reversed the effect on phase distribution, proliferation, mitochondrial stress, and *TOMM7* expression. Finally, through RNA-seq analysis we identified several significantly opposite expressed genes in CRISPRa compared to CRISPRi as *MYC*-responsive genes enriched for GO terms that included cell cycle G1/S transition, G2/M checkpoints, and DNA replication. *SNHG26* is itself a direct *MYC* target gene, and our results suggest that it affects cell cycle and proliferation through feed forward regulation of other *MYC*-responsive genes.

## Results and discussion

### *SNHG26* is mainly localized in the nucleus

*SNHG26* is localized at chromosome 7 (ENSG00000228649, chr7:22,854,126-22,872,945) between protein coding genes *TOMM7* and *FAM126A* (Figure 1A). Based on tissue expression data from the Genotype-Tissue Expression (GTEx) project, *SNHG26* is expressed across most tissue types, except for the brain, where it is barely detected (Figure 1A; Supplementary Figure S1). The primary sequence of *SNHG26* has limited conservation outside primates, but the *TOMM7*-*FAM126A* locus is conserved in jawed vertebrates and the snoRNA *SNORD93*, located in *SNHG26*’s second intron, is conserved in amniota (Supplementary Figure S2) (Gardner et al., 2015). Moreover, the genomic blocks containing *SNHG26*’s exons are largely conserved among placental mammals, and the genomic region surrounding *SNORD93* in both placentals and other amniota is transcribed (Supplementary Figure S3). Thus, *SNHG26* appears to have both conserved exon-intron structure, synteny, and expression in placentals.

**Figure 1.**
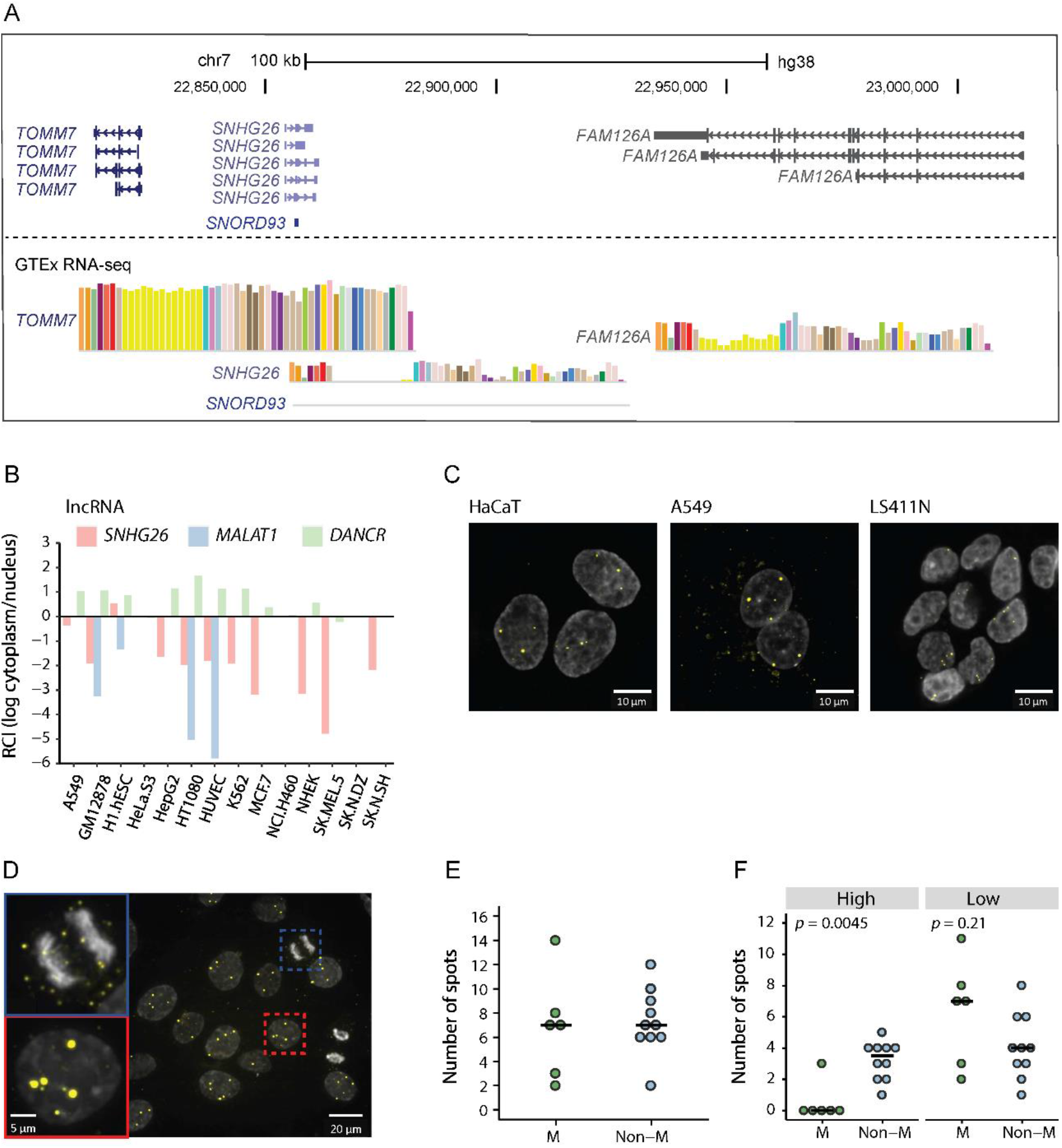
*SNHG26* is mainly localized in the nucleus as bright foci, and as nuclear and cytoplasmic low-intensity spots. (A) The genomic loci of SNHG26 from the UCSC Genome Browser (https://genome.ucsc.edu/). Tissue expression data (transcript per kilobase million, TPM) are from the GTEx project. (B) The subcellular localization of *SNHG26, MALAT1*, and *DANCR*. Data are from the lncAtlas (https://lncatlas.crg.eu/) and display the subcellular localization based on the RCI of RNA between cytoplasm and nucleus. Values < 0 indicate nuclear enrichment. (C) The localization of *SNHG26* (yellow) in fixed HaCaT, A549, and LS411N cells with DAPI-stained nuclei (grey) were visualized by using SNHG26-specific Stellaris™ probe sets conjugated to a Quasar670 dye in the 3′ end. Presented images are maximum intensity projections of 34 Z-stack slices (7.26 μm) of the cell. (D) A representative image of mitotic HaCaT cells. (E) The total number of spots co-localized with the DNA in 6 mitotic (M) and 10 non-mitotic (non-M) HaCaT cells. Crossbars represent median values. For the spot intensity analysis, a circular ROI of 10 pixels was used for manually measuring the intensity of individual spots that were associated with the DNA and a cut-off of 5000 was used to differentiate between high- and low-intensity spots. A Wilcoxon rank sum test was used to compare the distribution in mitotic and non-mitotic cells (*p*-value = 0.869). (F) The total number of high- and low-intensity spots co-localized with DNA in 6 mitotic (M) and 10 non-mitotic (non-M) HaCaT cells. Crossbars represent median values. A Wilcoxon rank sum test was used to compare the distribution in mitotic and non-mitotic cells; high-intensity spots *p*-value = 0.0045, low-intensity spots *p*-value = 0.208.

Data from the lncATLAS database (https://lncatlas.crg.eu/), which present nuclear localization as the log ratio of relative nuclear to cytoplasmic RNA levels as measured by RNA-seq (relative concentration index, RCI), indicate that *SNHG26* is enriched in the nucleus across a broad set of cell lines (average RCI = −2.03, Figure 1B). In comparison, the nuclear and cytoplasmic lncRNAs *MALAT1* and *DANCR* have average RCIs of −5.12 and 0.68, respectively (Figure 1B). We confirmed the subcellular localization of *SNHG26* using RNA FISH in HaCaT, A549, and LS411N cells. There, *SNHG26* is mainly localized in the nucleus where it appears as two to five bright nuclear foci as well as nuclear and extranuclear low-intensity spots (Figure 1C). The appearance of *SNHG26* as low and high intensity spots may represent *trans* and *cis* functional mechanisms, respectively.

The bright nuclear foci are consistent with functions connected to chromatin regulation and imprinting, such as for *XIST* and *AIR* [8, 16]. These lncRNAs accumulate at their site of action, and their presence suggests active silencing of the specific genomic location (Wang and Chang, 2011). Bright foci may also be consistent with the appearance of lncRNAs that are localized in subnuclear compartments or structures, such as Cajal bodies, the nucleolus, paraspeckles, nuclear speckles, perinucleolar, and perichromatin regions (Singh and Prasanth, 2013, Fay and Anderson, 2018). As HaCaT, LS411N, and A549 are all characterized as aneuploid cell lines with chromosome numbers that may be compatible with the number of high-intensity spots observed in these cell lines (Knutsen et al., 2010, Briffa et al., 2015, Peng et al., 2010, Boukamp et al., 1988), the high-intensity spots of *SNHG26* in the nucleus are possibly caused by an accumulation of its transcripts at their site of transcription.

According to Cabili *et.al*., the appearance of bright nuclear foci could imply that the lncRNA is involved in maintaining the epigenetic state during cell division, and to address this they investigated whether the nuclear foci persisted and remained attached to the chromosomes through mitosis (Cabili et al., 2015). To examine whether the nuclear high-intensity spots of *SNHG26* would persist during mitosis, we compared the intensity and number of spots in mitotic and non-mitotic HaCaT cells. The average number of spots associated with the DNA was similar in mitotic (6.8) and non-mitotic (7.3) cells (Figure 1D and 1E), yet few high-intensity spots (>5000 intensity) were detected in cells undergoing mitosis compared to non-mitotic cells (Figure 1D and 1F; Supplementary Figure S4). Spots associated with the DNA in mitotic and non-mitotic cells had an average intensity of 1495 and 8015, respectively. Based on these results, high-intensity spots are largely lost in mitotic cells, yet the number of spots that co-localize with DNA is similar. Thus, we cannot exclude that *SNHG26* may have functions connected to epigenetic maintenance through cell division, which warrants further investigations.

We only included the spots that co-localized with the stained DNA for the image analysis (Figure 1E-F), which excluded spots that were localized outside the DNA in mitotic cells, yet possibly within the nuclear membrane (Supplementary Figure S5). Consequently, the total number of spots present within the nucleus are probably higher in mitotic cells than the analysis suggests. This was supported by a visual inspection, where it appears to be several low-intensity spots surrounding the DNA in mitotic cells, probably within the nuclear membrane (Supplementary Figure S5).

### *SNHG26* affects proliferation and cell cycle phase distribution

Our previous data showed that knockdown of *SNHG26* resulted in growth inhibition in four different cell lines (preprint: https://doi.org/10.1101/2021.02.12.430890) 48 hours after transfection. To investigate whether this effect was consistent over time, we used ASO to knockdown *SNHG26* in HaCaT, A549, and LS411N cells (Supplementary Figure S6), followed by cell counting. A significant growth reduction of cells was observed in all three cell lines, with an average reduction of 32% and 57%, 48 and 72 hours after transfection, respectively (Figure 2A).

**Figure 2.**
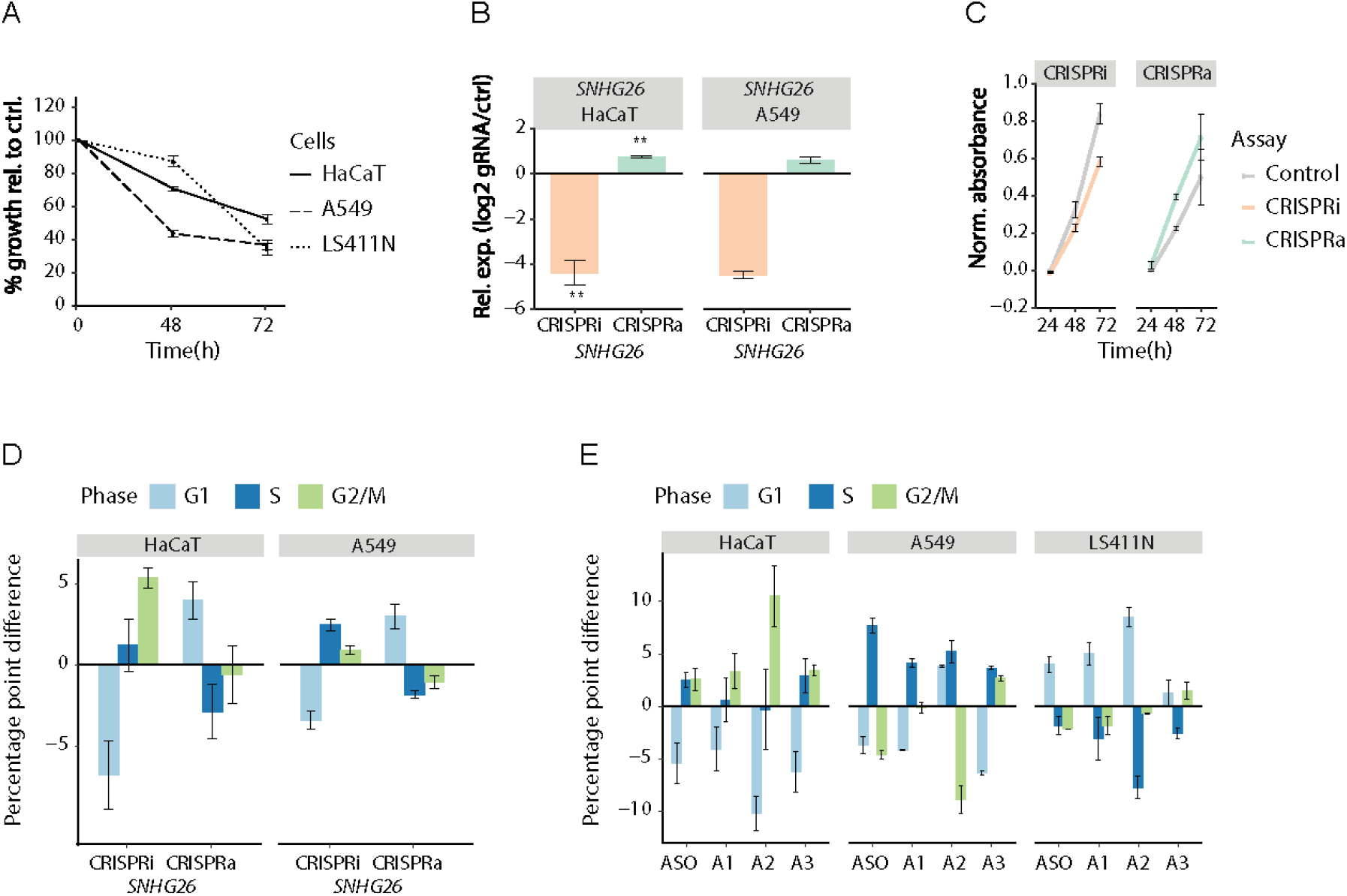
*SNHG26* affects proliferation, metabolic activity, and cell cycle phase distribution. (A) Effect of ASO-mediated knockdown of *SNHG26* on proliferation in HaCaT, A549, and LS411N cells. Data are the number of cells following ASO treatment relative to control treated cells (percentage of control) as measured by cell counting. Bars and error bars are mean and standard error of mean (SEM) of three or more independent replicates. Significant differences were determined by Student’s t-test (unpaired, two-tailed), assuming equal variances (all *p*-values \*\**p* ≤ 0.01), except LS411N 48 h with *p* = 0.058)). (B) The relative expression level of *SNHG26* as measured by RT-qPCR in response to CRISPRi/a of *SNHG26* in HaCaT and A549 cells. Data are presented as fold change expressions of cells transduced with *SNHG26*-specific gRNA relative to control gRNA. Bars and error bars are mean and SEM of two or more independent replicates. Significant differences were determined by Student’s t-test (unpaired, two-tailed), assuming equal variances (\**p* ≤ 0.05; \*\**p* ≤ 0.01). (C) Effect of CRISPRi/a of *SNHG26* on metabolic activity in HaCaT cells as measured by XTT. Data are presented as normalized absorbance (A_465 nm_ − A_630 nm_) for cells transduced with *SNHG26*-specific gRNA and control gRNA. Bars and error bars are mean and SEM of three independent replicates. Significant differences were determined by Student’s t-test (paired, two-tailed), assuming equal variances (CRISPRi 48 h *p* = 0.167, 72 h *p* = 0.061; CRISPRa 48 h *p* = 0.004, 72 h *p* = 0.027). (D-E) The distribution of cells in G1, S, and G2/M cell cycle phases in response to (D) CRISPRi/a modulation of *SNHG26* in HaCaT and A549 cells and (E) ASO and siRNA (A1, A2, A3) mediated knockdown of *SNHG26* in HaCaT, A549, and LS411N cells. Data are the difference in percentages of G1, S, and G2/M cells between cells transduced/transfected with target-specific gRNA/siRNA/ASO to those transduced/transfected with a control gRNA/siRNA/ASO. (D) Bars and error bars are mean and SEM of three independent replicates. ANOVA *p*-values were calculated from a hierarchical, linear model: HaCaT: G1: 7e-04, S: 0.120, and G2/M: 0.031; A549: G1: 5.02e-08, S: 5.83e-06, and G2/M: 0.007. (E) Bars and error bars are mean and SEM of two or more independent replicates. ANOVA *p*-values were calculated from a hierarchical, linear model: HaCaT : G1: 7.96e-06, S: 0.275, and G2/M: 0.0002; A549 : G1: 0.068, S: 0.001, and G2/M: 0.056; LS4111N: G1: 6.05e-05, S: 0.0005, and G2/M: 0.387.

To further investigate the functions and biological effects of *SNHG26* expression, we generated HaCaT and A549 cells stably expressing catalytically inactive CAS9, dead CAS9 (dCAS9), fused with either the Krüppel-associated box (KRAB) repressing domain or multiple activating domains, providing tools for, respectively, transcriptional inhibition (CRISPRi) or activation (CRISPRa) (Figure 2B; Supplementary Figure S7). Consistent with the reduced cell counts following ASO and siRNA-mediated knockdown of *SNHG26*, transcriptional inhibition of *SNHG26* reduced the metabolic activity of HaCaT cells, while its activation increased the metabolic activity (Figure 2C). The reduction of metabolic activity was validated by siRNA-mediated knockdown of *SNHG26* in HaCaT cells, with 12% reduction 48 hours after transfection (Supplementary Figure S8).

Further we used CRISPRi/a to investigate whether modulation of *SNHG26* transcription affected the cell cycle distribution in HaCaT and A549 cells. We used a hierarchical, linear model to calculate the combined effect on cell cycle phase distribution between CRISPRi and CRISPRa, requiring opposite contributions of the two assays and assuming a random effect for each assay. In HaCaT cells we observed a significant difference in cell cycle distribution for the G1 and G2/M phases, whereas for A549 we observed a significant difference for all cell cycle phases (G1, S, and G2/M) between CRISPRi and CRISPRa (Figure 2D). For both cell lines, CRISPRi resulted in a reduction of cells in the G1 phase and an enrichment of cells in the G2/M phase, though for A549 cells, the enrichment in the S phase was higher (Figure 2D). As expected, the opposite trend was observed when upregulating *SNHG26* by CRISPRa, which resulted in an enrichment and reduction in the G1 and S phases, respectively, of both cell lines (Figure 2D). To investigate whether the cell cycle phase distribution was stable over time, CRISPRi/a-modified HaCaT cells were harvested and re-seeded followed by cell cycle assays eight days after gRNA transduction with *SNHG26*-specific gRNA or non-target control gRNA. Indeed, the cell cycle phase distribution was the same as before re-seeding with significant difference in cell cycle distribution for the G1 and G2/M phases between CRISPRi and CRISPRa, suggesting that the effect is not transient, but stable over time (Supplementary Figure S9).

To investigate whether the effect on cell cycle phase distribution was consistent when using other gene modulation techniques, we performed ASO- and siRNA-mediated knockdown of *SNHG26* in HaCaT, A549, and LS411N cells (Supplementary Figure S6). For each cell line, we used a hierarchical, linear model to calculate the effect on cell cycle phase distribution across the four different ASO/siRNA assays assuming a random effect for each assay. In line with previous results, we observed a significant difference in cell cycle distribution for the G1 and G2/M phases in HaCaT cells. A549 cells showed a significant difference in cell cycle distribution for the S phase, whereas for LS411N cells, the cell cycle distribution was significant for the G1 and S phases (Figure 2E).

While the siRNAs and ASO used affected cell cycle distribution in A549 cells, the phase distribution varied across assays, probably due to assay-specific off-target effects. The number of cells present in the G1 phase was reduced using ASO and in two out of three siRNAs, while there was a significant enrichment of cells in the S phase across all four assays. Meanwhile the effect on the G2/M phase varied, with a reduction of cells present in the G2/M phase using ASO and siRNA A2, no effect with siRNA A1, and an increase with siRNA A3 (Figure 2E).

The effect of knockdown was more consistent between different siRNAs and the ASO in LS411N cells. Interestingly, ASO and siRNA-mediated knockdown of *SNHG26* had the opposite effect on phase distribution in LS411N cells as compared to what we observed in HaCaT and A549 cells, with a significant enrichment of cells in the G1 phase and a reduction of cells in the S phase across all four assays (Figure 2E). This implies that *SNHG26* affects cell cycle through a different mechanism in LS411N, possibly related to cell-specific mechanisms and differences in the cells’ epigenetic or mutational status.

Importantly, the effects of modulating *SNHG26* expression were consistent between the different methods used. Unintended effects caused by non-specific binding are more or less frequent for siRNAs, ASO, and CRISPRi/a, but differ between methods (Stojic et al., 2018, Jackson et al., 2003). Such method-specific biases and off-target effects are therefore unlikely explanations of our results.

### *SNHG26* regulates the expression of its neighbor gene TOMM7

The mainly nuclear localization pattern as well as the appearance of bright foci we observed for *SNHG26* may be consistent with an accumulation of transcripts at its site of transcription. Thus, we wondered whether *SNHG26* regulates the expression of other genes from its location of transcription. To address this question, we used RT-qPCR to determine whether CRISPRi/a of *SNHG26* affected the expression level of its neighbor mRNAs *TOMM7* and *FAM126A* in HaCaT cells. We observed a reduction in the expression of *TOMM7* using CRISPRi, and the opposite trend for CRISPRa. Meanwhile, CRISPRi did not affect the expression of *FAM126A*, whereas CRISPRa of *SNHG26* increased the expression of *FAM126A*, although not significantly, due to large variation between replicates (Figure 3A).

**Figure 3.**
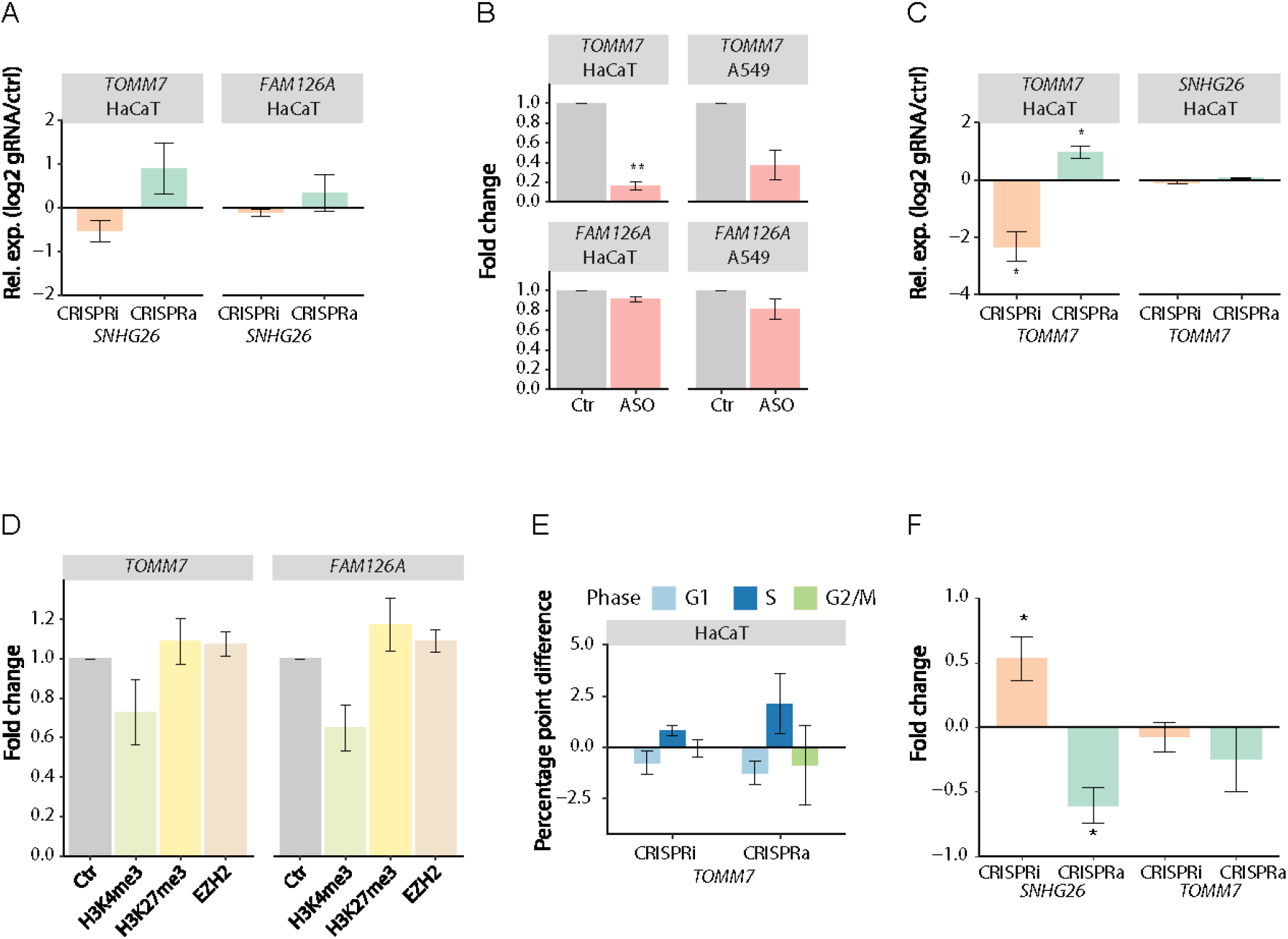
SNHG26 affects the expression of neighbor genes and mitochondrial stress. (A) The relative expression levels of *TOMM7* and *FAM126* as measured by RT-qPCR in response to CRISPRi/a of *SNHG26* in HaCaT cells. Data are presented as fold change expressions of cells transduced with target-specific gRNA relative to control gRNA. Bars and error bars are mean and SEM of three independent replicates (n = 2 for CRISPRa FAM126A). *p*-values were calculated by Student’s t-test, (unpaired, two-tailed) assuming equal variances (*TOMM7 p* = 0.06 and *p* = 0.175; *FAM126 p* = 0.24 and *p* = 0.47 for CRISPRi and CRISPRa, respectively). (B) The relative expression of *TOMM7* and *FAM126* as measured by RT-qPCR in HaCaT and A549 cells. Data are presented as fold change expressions of target genes following ASO treatment relative to control-treated cells. Bars and error bars are mean and SEM of three independent replicates. Significant differences were determined by Student’s t-test, (unpaired, two-tailed) assuming equal variances (\*\**p* ≤ 0.01, *TOMM7 p* = 0.003 and *p* = 0.052; *FAM126 p* = 0.085 and *p* = 0.212 in HaCaT and A549, respectively). (C) The relative expression level of *TOMM7* and *SNHG26* as measured by RT-qPCR in response to CRISPRi/a of *TOMM7* in HaCaT cells. Data are presented as fold change expressions of cells transduced with target-specific gRNA relative to control gRNA. Bars and error bars are mean and SEM of three independent replicates. Significant differences were determined by Student’s t-test, (unpaired, two-tailed) assuming equal variances (\**p* ≤ 0.05). (D) We used ChIP-qPCR to evaluate the effect of CRISPRi of *SNHG26* on the level of H3K4me3, H3K27me3, and EZH2 at the TSS of *TOMM7* and FAM126A in HaCaT cells. Bars and error bars are mean and SEM of three independent replicates. *p*-values were calculated by Student’s t-test, (unpaired, two-tailed) assuming equal variances (*TOMM7 p* = 0.245, *p* = 0.519, *p* = 0.338; *FAM126 p* = 0.09, *p* = 0.326, *p* = 0.243, for H3K4me3, H3K27me3, and EZH2, respectively). (E) The distribution of cells in G1, S, and G2/M cell cycle phases in response to CRISPRi/a of *TOMM7* in HaCaT cells. Data are the difference in percentages of G1, S, and G2/M cells between cells transduced with target-specific gRNA to those transduced with a control gRNA. Bars and error bars are mean and SEM of three independent replicates. (F) We used Mitosox and FACS to determine whether CRISPRi/a-mediated modulation of *SNHG26* and TOMM7 expression affected mitochondrial stress in HaCaT cells. Data are presented as fold change fluorescence intensity of cells transduced with target-specific gRNA relative to control gRNA. Bars and error bars are mean and SEM of three or more independent replicates. Significant differences were determined by Student’s t-test (unpaired, two-tailed), assuming equal variances (\**p* ≤ 0.05).

To address whether *SNHG26*’s effect on its neighboring genes was transcriptional or post-transcriptional, we first investigated *TOMM7* and *FAM126A* expression following ASO-mediated knockdown of *SNHG26*. In both HaCaT and A549 cells, we observed a downregulation of *TOMM7* expression (Figure 3B). We also observed a slight downregulation of *FAM126A* in response to ASO-mediated knockdown in both cell lines, yet the effect was not significant. Second, we investigated whether CRISPRi/a of *TOMM7* affected *SNHG26* expression, reasoning that if *SNHG26*’s effect was primarily post-transcriptional we should see no effect on *SNHG26* expression. Indeed, CRISPRi/a mediated knockdown and upregulation of *TOMM7* had no effect on the expression of *SNHG26* (Figure 3C).

We further investigated whether downregulation of *SNHG26* affected the chromatin state at the TSSs of *TOMM7* and *FAM126A*. We used CRISPRi to knockdown *SNHG26* in HaCaT cells followed by ChIP-qPCR of H3K4me3, H3K27me3, and EZH2. EZH2 was included as an additional marker for gene repression, as it is a part of the Polycomb Repressive Complex 2 (PRC2) where it catalyzes the addition of methyl groups to lysine 27 of histone 3 (Cao et al., 2002). Consistent with reduced transcriptional activity, we observed a reduction of H3K4me3 occupancy at the TSS of both *TOMM7* and *FAM126A,* and a slight increase in H3K27me3 and EZH2, yet none were significant (Figure 3D).

Overall, the above results suggest that *SNHG26* is a post-transcriptional positive regulator of *TOMM7*, possibly through *cis* accumulation of *SNHG26* RNA.

### *SNHG26* affects mitochondrial stress

TOMM7 is a part of the TOM complex found in the mitochondrial membrane where it is involved in the transport of proteins into the mitochondria. TOMM7 probably acts by stabilizing PTEN-induced kinase 1 (PINK1) on the outer membrane of depolarized mitochondria where it recruits Parkin and activates its E3 ubiquitin ligase by phosphorylation. Impaired stabilization of PINK1 results in less activation of Parkin, which is necessary for the degradation of damaged mitochondria through mitophagy, and causes increased production of reactive oxygen species (ROS) and cell death (Pickrell and Youle, 2015). Due to the essential roles of mitochondria in cellular homeostasis, mitochondrial damage is associated with a broad spectrum of diseases (Palikaras et al., 2018). Results from a genome wide RNAi screen suggest that loss of *TOMM7* blocks the accumulation of PINK1 on the outer mitochondrial membrane (OMM) and the recruitment of PARKIN in response to chemically depolarized OMM, yet the import of proteins through the TOM complex is not affected (Hasson et al., 2013, Sekine et al., 2019). Previous studies have shown that TOMM7 is a driver of cerebral angiogenesis, and *TOMM7* deficiency affects mitochondrial stress in primary cultures of human brain microvascular endothelial cells and in vascular-specific transgenic zebrafish (Shi et al., 2018).

As *SNHG26* regulates the expression of *TOMM7*, we sought out to examine whether *SNHG26* induced growth inhibition was caused by the downregulation of *TOMM7*, due to increased ROS production and subsequent cell death. To address this question, we measured the level of mitochondrial ROS in response to CRISPRi/a of *SNHG26* and *TOMM7* in HaCaT cells. CRISPRi-mediated downregulation of *SNHG26* resulted in a significant increase of mitochondrial stress, and we observed an opposite trend for the CRISPRa-mediated upregulation of *SNHG26* (Figure 3F). The increased mitochondrial stress after *SNHG26* downregulation was validated by ASO-mediated knockdown of *SNHG26* in HaCaT and A549 cells (Supplementary Figure S10). In contrast, there was no significant effect of CRISPRi/a-mediated modulation of *TOMM7* on the level of mitochondrial stress (Figure 3F). Our results show that increasing and decreasing *SNHG26* expression affect *TOMM7* expression and mitochondrial stress, but that directly modulating *TOMM7* expression has no effect on mitochondrial stress. Consequently, *SNHG26* affects mitochondrial stress through a mechanism that is either independent off or synergistic with *SNHG26*’s effect on *TOMM7* expression.

### RNA sequencing identifies differentially expressed genes involved in cell cycle, DNA replication, and proliferation

Whereas the above results showed that *SNHG26* is a positive regulator of its neighboring gene *TOMM7*, the results also suggested that *SNHG26* has additional *trans* effects. Specifically, as *SNHG26* is localized both as bright nuclear foci and as low-intensity spots, we reasoned that *SNHG26* probably impacts diverse target genes post-transcriptionally by relocating or acting in *trans* from its site of transcription.

To identify such possible *trans* targets, we performed RNA-seq of CRISPRi and CRISPRa in HaCaT cell lines transduced with *SNHG26*-specific gRNA or a non-target control gRNA (Figure 4A). For CRISPRi and CRISPRa we identified a total of 1692 and 1268 differentially expressed genes with *p*-value ≤ 0.05, respectively. Moreover, there were 110 differentially expressed genes with *p*-value <0.05 in both CRISPRi vs. Control and CRISPRa vs. Control (Supplementary Figure S11). Of these, 61 genes were differential expressed in opposite directions in the CRISPRi compared with the CRISPRa experiments (*p*-value ≤ 0.01), including *LATS1* (large tumor suppressor 1) and *LIN52* (protein lin-52 homolog).

**Figure 4.**
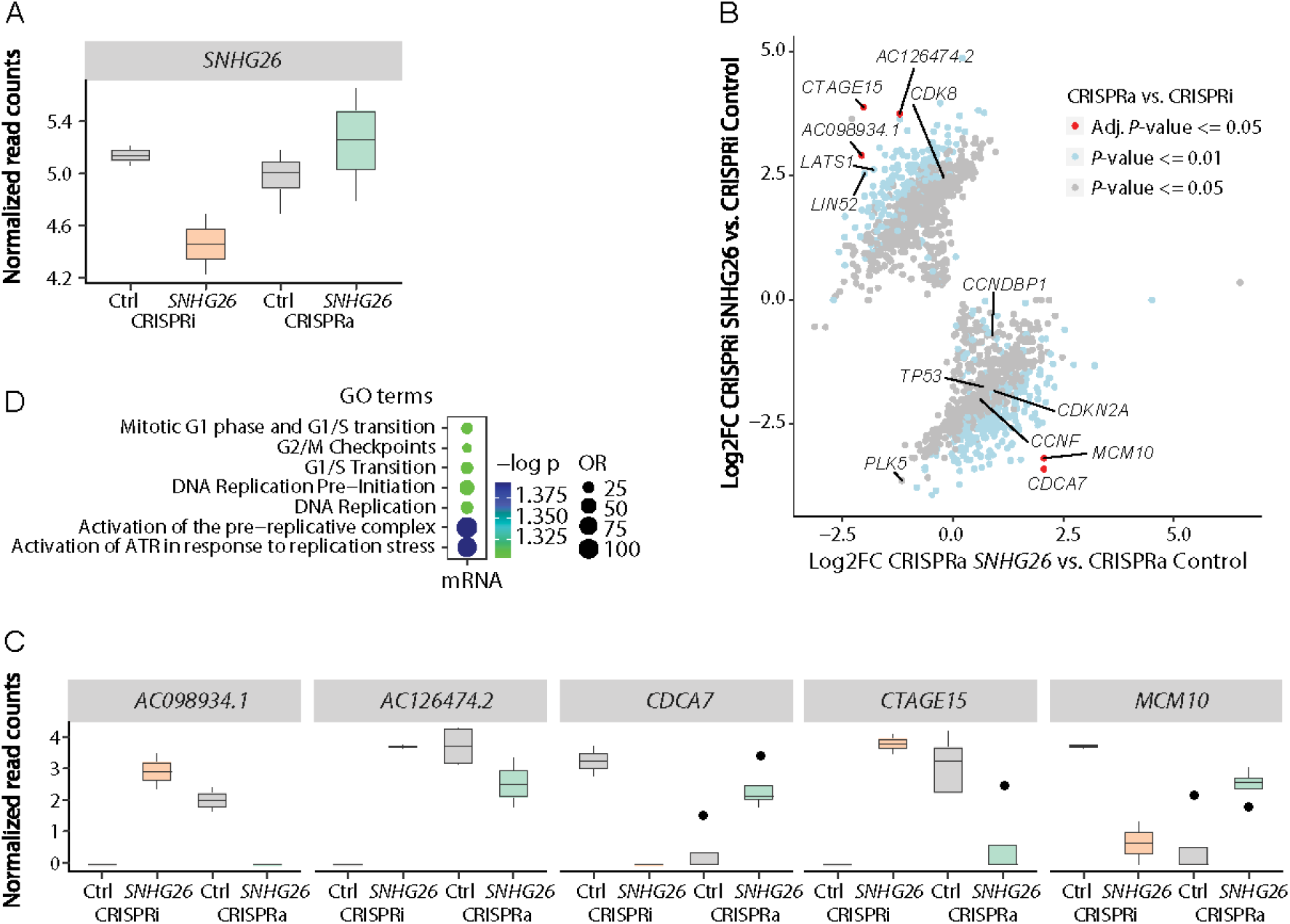
Total RNA-seq of CRISPRi/a modified HaCaT cell lines identify differentially expressed genes with functions connected to cell cycle and proliferation. (A) RNA-seq expression level of *SNHG26* in CRISPRi (n=2) and CRISPRa (n=4) modified HaCaT cells transduced with a *SNHG26*-specific gRNA compared to a non-specific gRNA control. Data are presented as normalized read counts. (B) A total of 1623 genes were differentially affected in CRISPRa vs. CRISPRi with *p*-value < 0.05. They are colored according to their *p*-values and interesting genes are labelled. (C) The expression of the top five significant genes that were differentially expressed in the opposite direction in CRISPRa compared to CRISPRi. Data are presented as normalized read counts with adjusted *p*-value < 0.05. (D) GO enrichment analysis for the top five significant genes that were differentially expressed in the opposite direction in CRISPRa compared to CRISPRi. The results show enrichment of cell cycle related GO terms.

As we expected targets of *SNHG26* to be differentially affected in CRISPRi compared with CRISPRa of *SNHG26*, we did a separate interaction analysis to identify such genes. This analysis identified 1623 genes that were differentially expressed between CRISPRa vs. Control compared to CRISPRi vs. Control with *p*-value ≤ 0.05, which included well-known cell cycle regulators such as *TP53, CCNF* (cyclin F)*, CDK8, CDKN2A* (cyclin-dependent kinase inhibitor 2A), *PLK5* (polo like kinase 5), and *CCNDBP1* (cyclin D1 binding protein 1) (Figure 4B; Supplementary Dataset S1). The top five significant genes (adjusted *p*-value ≤ 0.05, Figure 4C) were significantly enriched for GO terms related to cell cycle, including G1/S transition, G2/M checkpoints, and DNA replication (Figure 4D), which is in line with our data that demonstrates that *SNHG26* affects cell cycle progression and proliferation.

Three of the five genes, *MCM10* (minichromosome maintenance protein 10), *CDCA7* (cell division cycle associated 7), and *AC098934.1*, have previously been described in the literature with growth promoting abilities (Gao et al., 2020, Park et al., 2008, Osthus et al., 2005). In our study, *MCM10* was downregulated (log_2_ fold change (LogFC −3.185) in CRISPRi cells transduced with gRNA targeting *SNHG26* and upregulated (LogFC 2.058) in response to CRISPRa-mediated upregulation of *SNHG26* compared to control. Another growth promoting protein included in the top five opposite expressed genes was *CDCA7*, which was also downregulated (LogFC −3.403) in response to CRISPRi and upregulated (LogFC 2.059) in response to CRISPRa of *SNHG26*. The lncRNA *AC098934.1* (also known as *RP11-480I12.5* and *TUBAP5*) was upregulated (LogFC in 2.909) in response to CRISPRi and downregulated (LogFC in −2.054) in response to CRISPRa of *SNHG26.* Meanwhile, *AC126474.2* and *CTAGE15* have unknown functions.

Several of the differentially expressed genes in both CRISPRi and CRISPRa are direct or indirect MYC responsive genes, including *MCM10, CDCA7, TP53, CDKN2A, CCNF*, and *CDK8* (Yang et al., 2018, Prescott et al., 2001, Koch et al., 2007, Kang et al., 2020, Gill et al., 2013, Kidder et al., 2008, Phesse et al., 2014, Garcia-Gutierrez et al., 2019). *SNHG26* was detected as a MYC-target in P493-6 human B-cells, where it was upregulated in response to *MYC* overexpression (Hart et al., 2014). A subsequent study found *SNHG26* to be a direct transcriptional target of *MYC* and a driver of *MYC*-mediated proliferation in human lymphoid cell lines (Raffeiner et al., 2020). From our previous study on cell cycle synchronized HaCaT cells, the level of *MYC* in HaCaT cells peaked during the G2/M phase, similarly to the cell cycle profile of *SNHG26,* demonstrating a positive correlation between their expression through the cell cycle phases (preprint: https://doi.org/10.1101/2021.02.12.430890). The *MYC* family of oncogenes consists of *c-MYC, N-MYC*, and *L-MYC*, and is believed to regulate the transcription of about 15% of the genome. *MYC* regulates several of the positive cell cycle regulators including cyclins, CDKs and E2F transcription factors. Moreover, *MYC* is able to inhibit the activation of the cell cycle inhibitors p21 and p27, and is overexpressed in 60-70% of human cancers (Garcia-Gutierrez et al., 2019). Studies suggest cell cycle-associated functions of c-MYC for different cancer cells, including regulatory roles for the G1/S and G2/M phase transitions through different mechanisms (Felsher et al., 2000, Song et al., 2013, Yang et al., 2018).

Both *MCM10* and *CDCA7* were identified as cell cycle-associated genes in our previous study, with expression levels peaking during G1/S phase (preprint: https://doi.org/10.1101/2021.02.12.430890). MCM10 is a known cell cycle-associated growth promoting protein that is required for proper DNA replication (Baxley and Bielinsky, 2017). MCM10 plays an essential role in initiation of DNA replication at replication origins and elongation by stabilizing DNA polymerase-α and its association to chromatin during DNA replication. Downregulation of MCM10 reduces the replication fork elongation rate. Moreover, MCM10 is a CDK substrate and is associated with chromatin in the late stage of replication initiation, although it is not necessary for the process (Chadha et al., 2016). *MCM10* is often overexpressed in several cancers, and its expression is associated with tumor progression (Kang et al., 2020, Cui et al., 2018, Baxley and Bielinsky, 2017)). Moreover, *MCM10* is listed as one of the top ranked cancer-associated genes in non-small-cell lung cancer (NSCLC) and breast cancer across different methods and from independent patient datasets (Wu et al., 2012). Depletion of *MCM10* has been reported to induce G2/M cell cycle arrest (Park et al., 2008, Romani et al., 2015). A study identified *MCM10* as upregulated by *MYCN* in neuroblastoma, where it stimulated cell cycle progression. Although the expression levels of *MCM10* and *MYCN* were highly correlated, there were no direct interaction between *MYCN* and the promoter of *MCM10*, suggesting that *MCM10* is not a direct target of *MYCN*, or that *MYCN* binds outside the analyzed promoter region (Koppen et al., 2007). *CDCA7* is frequently upregulated in human cancers, including breast- and colorectal cancer, where its expression may be used to predict prognosis and progression (Li et al., 2020, Ye et al., 2018). Moreover, *CDCA7* is an E2F- and MYC responsive gene, and studies suggest that it contributes to MYC-mediated tumorigenesis, possibly by the direct association to MYC (Gill et al., 2013, Prescott et al., 2001, Osthus et al., 2005, Goto et al., 2006).

Our results show that *SNHG26* is predominantly nuclear and affects gene expression both in *cis* and *trans* – characteristics shared by other lncRNAs involved in transcriptional regulation (Cabili et al., 2015). However, since *SNHG26* also appears as extranuclear low-intensity spots, functions connected to the cytoplasm, such as *SNHG26* encoding a small peptide, cannot be excluded. We also note that *SNHG26* is host to snoRNA *SNORD93*, so CRISPR-mediated modulation of *SNHG26* expression could also affect *SNORD93*’s levels and thereby its targets. While it is less likely that siRNA- and ASO-mediated downregulation of *SNHG26* affects *SNORD93* expression, we cannot exclude this possibility.

In conclusion, downregulation of *SNHG26* affects several cellular processes including cell cycle, metabolic rate, proliferation, mitochondrial stress, and gene expression in *cis*- and in *trans,* an effect that appears to be post-transcriptional and may be reversed by upregulation of *SNHG26* expression. As *SNHG26* is a direct MYC target gene, we propose that it affects cell cycle and proliferation through the regulation of downstream MYC-responsive genes, such as *MCM10, CDCA7, TP53, CDKN2A, CCNF*, and *CDK8*

## Material and Methods

### Cell culture

All cell lines used were obtained from the American Type Culture Collection (ATCC) and cultivated in a humidified incubator at 37°C in 5% CO_2_. HaCaT, a human keratinocyte cell line, A549, an adenocarcinoma human alveolar basal epithelial cell line, and Hek293T, a human epithelial kidney cell line, were all cultured in Dulbecco’s modified Eagle’s medium (DMEM) (Sigma-Aldrich, D6429) supplemented with 10% fetal bovine serum (FBS) (Sigma-Aldrich, F7524) and 2 mM L-Glutamine (Sigma-Aldrich, G7513). For XTT viability and Mitosox assays we used DMEM without phenol red (Gibco, 21063-029). LS411N, a Dukes’ type B colorectal carcinoma cell line, was cultivated in RPMI 1640 Medium (Gibco, A1049101) supplemented with 10% FBS. All cell lines were sub-cultivated at least twice a week at about 70% confluence, and regularly tested for mycoplasma infections.

### Single molecule RNA fluorescence in situ hybridization (RNA FISH)

The software available through Stellaris Probedesigner was used to design the oligonucleotide sets. For the *SNHG26* we used masking 4, length 20 and spacing 2, which resulted in 37 oligonucleotides. Stellaris™ probesets (Biosearch Technologies) were conjugated to a Quasar670 dye in the 3′ end. Hybridization and staining were performed as prescribed in the Stellaris protocol for adherent cells. We used *GAPDH* as a predesigned control. The probe sequences are listed in Supplementary Table S1.

### Imaging

We used a Zeiss Laser TIRF 3 fluorescence microscope (Zeiss), equipped with a α-Plan-Apochromat 100x/1.46 oil-immersion objective for imaging of the cells. A Zeiss 81 HE DAPI/FITC/Rh/Cy5 filter was used, DAPI was excited by LED-module 365 nm (Zeiss Colibri) and Quasar670 was excited by 644 nm wavelength laser. The images were acquired by either a Hamamatsu EMCCD EMX2 or a Hamamatsu ORCA-Fusion camera at 16 bit and at a voxel size of 100 × 100 × 220 nm^3 (EMCCD) or 129 × 129 × 220 nm size (ORCA-Fusion). Image deconvolution was performed using SVI Huygenes Professional (version 18.10) and image analysis was done using Fiji (version 1.52t) (Schindelin et al., 2012). For the spot intensity analysis a circular region of interest (ROI) of 10 pixels was used for manually measuring the intensity of individual spots and a cut-off of 5000 was used to differentiate between bright and dim spots. Presented images are maximum intensity projections of 34 Z-stack slices (7.26 μm) of the cell.

### Plasmids and cloning

Lenti-sgRNA puro was a gift from Brett Stringer (Addgene plasmid # 104990). We followed the Zhang labs protocol (https://media.addgene.org/data/plasmids/52/52961/52961-attachment_B3xTwla0bkYD.pdf) for guide RNA (gRNA) design and cloning of the gRNA between the two BsmBI restriction sites. The hU6-F (5’-GAGGGCCTATTTCCCATGATT-3’) was used to sequence the RNA to validate gRNA insert. For CRISPRi we used pHR-SFFV-dCas9-BFP-KRAB (Gilbert et al., 2013), a gift from Stanley Qi & Jonathan Weissman (Addgene plasmid # 46911). For CRISPRa we used pXPR_120 with multiple activating domains VP64, P65 and Rta (Najm et al., 2018), a gift from John Doench & David Root (Addgene plasmid # 96917).

### Lentiviral production

Hek293T cells were plated 24 hours before transfection with Lenti-sgRNA puro containing the different gRNAs or pHR-SFFV-dCas9-BFP-KRAB or pXPR_120 together with the packaging plasmids psPAX.2 and pMD2.G using Lipofectamine 2000 (Invitrogen™, 11668019). Culture medium was replaced 7 hours after transfection to decrease toxicity. Fifty-five hours after transfection the viral supernatant was collected, centrifuged at 1800 g for 5 min at 4°C, filtered through a 45 μM filter, and stored at −80°C.

### Generating stable dCas9 expressing cell lines

We performed a reverse transduction of both HaCaT and A549 cells by adding lentiviral pHR-SFFV-dCas9-BFP-KRAB or pXPR_120 together with 8 μg/ml polybrene (Sigma-Aldrich, H9268) in DMEM, to generate CRISPRi and CRISPRa cell lines, respectively. The medium was replaced 48 hours after transduction, and 10 μg/ml blasticidin (Invivogen) was supplied to select for cells that had incorporated pXPR_120 into their genome. We changed the medium of pXPR_120 after three days, while selection was continued until all control cells had died, which took about five days. We kept the selection pressure on cells containing the activating construct pXPR_120 by supplying DMEM with 5 μg/ml blasticidin during sub-cultivating. We used a FACS Aria II cell sorter (BD Bioscience) to select for cells transduced with the inhibiting construct pHR-SFFV-dCas9-BFP-KRAB.

### Guide RNA (gRNA) design

The E-CRISPR online tool (http://www.e-crisp.org/E-CRISP) was used for gRNA design [12]. The sequence targeted −50 to + 200 base pairs (bp) relative to the transcriptional start site (TSS) of *SNHG26* (>hg38_dna range=chr7:22854048-22854298). We determined the exact location of TSS using the Fantom5 genome browser (ZENBU 3.0.1) (Lizio et al., 2015, Lizio et al., 2019). We used the Basic local alignment Search Tool to avoid off-target effects. The sequences of different gRNAs are listed in Supplementary Table S2.

### gRNA transductions

We transduced the CRISPRi/a modified cells with target-specific lentiviral gRNAs or a control gRNA together with 8 μg/ml polybrene 20 hours after seeding at approximately 50% confluence. The multiplicity of infection (MOI) was 2.1. To select for resistant cells that contained the gRNA, we added 2 μg/ml puromycin (Invivogen, ant-pr-1) to the growth media 24 hours after transduction. Cells were harvested 72 hours after selection started. We included gRNAs specific for *MALAT1* and *SLC4A1* as positive controls for CRISPRi and CRISPRa, respectively.

### Changes in cell cycle phase distribution are not caused by *TOMM7*

To investigate whether the observed changes in cell cycle distribution in response to CRISPRi/a of *SNHG26* is mediated through the regulation of *TOMM7,* we used CRISPRi/a to modulate the expression of *TOMM7* in HaCaT cells, followed by cell cycle assays (Figure 3E). We did not observe the same changes in cell cycle phase distribution as for *SNHG26* (Figure 2D), suggesting *SNHG26* is affecting the cell cycle through another mechanism.

### Short interfering (siRNA) and Antisense oligo (ASO)-mediated knockdown

We transfected cells with siRNAs and/or Antisense LNA GapmeRs (Antisense oligo; ASOs) for a 20 nM final concentration using Lipofectamine RNAimax (Invitrogen™, 13778030) when seeded, according to the manufacturer’s protocol. Cells were harvested after 48 and/or 72 hours after transfection at about 70% confluence. MISSION^®^ siRNA Universal Negative Control #1 (Sigma, SIC001) and Negative control B Antisense LNA GapmeR (Qiagen, LG00000001) were used as controls for siRNAs and ASOs, respectively. The producers and sequences of siRNAs and ASOs are listed in Supplementary Table S3.

### Quantitative Real-Time PCR (RT-qPCR)

To isolate total RNA, we used the *mir*Vana miRNA Isolation Kit (Invitrogen™, AM1560) according to manufacturer’s protocol, followed by DNase treatment with TURBO DNA-*free*™ (Invitrogen™, AM1907). We used NanoDrop ND-1000 UV-Vis Spectrophotometer to measure the RNA concentration and quality. RNA was converted to cDNA under standard conditions with random hexamer primers using TaqMan™ Reverse Transcription reagents (Invitrogen™, N8080234). Quantitative RT-PCR reactions were prepared with SYBR select master mix (Applied Biosystems, 4472919). The sequences of the different primers used are listed in Supplementary Table S4. Relative expression levels were calculated using the comparative C_T_ method (2^−ΔΔCT^ method (Livak and Schmittgen, 2001)), and expression data were normalized to *GAPDH*.

### Viability assay

To investigate how up- and downregulation of *SNHG26* affected viability, measured as the level of metabolic activity, we performed TACS XTT Cell viability Assay (R&D Systems™, 4891025K). HaCaT cells with the repressing domain and the activating domains were transduced with target-specific lentiviral gRNAs or a non-target control gRNA. Cells were seeded in triplicates for each condition in a 96-well tray 96 hours after transduction. We measured absorbance 24, 48, and 72 hours after seeding, 4 hours after XTT was added, following the manufacturer’s protocol. XTT viability assay was also performed with HaCaT cells transfected with *SNHG26*-specific siRNA or a non-target control siRNA as described in the section above. We measured absorbance 48 hours after seeding, 4 hours after XTT was added, following the manufacturer’s protocol.

### Proliferation assay

We performed cell counting using Moxi z mini automated cell counter (ORFLO Technologies) to investigate how ASO-mediated knockdown of *SNHG26* affected proliferation in HaCaT, A549, and LS411N cells. Cells were seeded in triplicates for each condition in a 24-well tray and counted 48 and 72 hours after transfection. Each well was washed twice with preheated PBS and trypsinated for 8-10 min before the cells were resuspended in preheated growth medium and counted. We applied a two-tailed, paired Student’s *t*-test to test whether the cell number was significantly different (*p-* value < 0.05) between cells transfected with a non-target negative control ASO and a target-specific ASO in three or more independent experiments.

### Cell cycle and fluorescence-activated cell sorting (FACS) analysis

For the cell cycle experiments cells were harvested 48 hours after siRNA-mediated transfection, and 96 hours after gRNA transduction. We washed twice with preheated PBS and trypsinated for 8-10 min before collecting the cells using ice-cold PBS supplemented with 3% FBS. The cells were then centrifuged at 4°C for 5 min, and the supernatant was carefully removed. For FACS analysis, cells were fixed in ice-cold 100% methanol and stored at 4°C until DNA measurement. The cells were washed with cold PBS and incubated with 200 μl of DNase-free RNAse A in PBS (100 μ g/ml) for 30 min at 37°C before DNA staining with 200 μl of Propidium Iodide (PI, Sigma) (50 μ g/ml) at 37°C for 30 min. We used BD FACS Canto flow cytometer (BD Biosciences) for cell cycle analyses. The excitation maximum of PI is 535 nm and the emission maximum is 617 nm. The blue laser (488 nm) excited PI-stained cells and the PI fluorescence was detected in the Phycoerythrin PE channel (578 nm). Cell cycle fractions were determined by using the FlowJo software.

### Chromatin Immunoprecipitation (ChIP)

HaCaT cells were trypsinated and harvested with PBS containing 3% FBS and washed twice with PBS (RT, room temperature). We counted the cells, and 3x 10^6^ cells were added to a final volume of 500 μl PBS. For crosslinking we added 13.5 μl formaldehyde for a final concentration of 1%, vortexed gently, and incubated for 10 minutes at RT. To stop the crosslinking we used 50 μl 1.25 M glycine for a final concentration of 0.125 M, vortexed gently and incubated for 5 min at RT. Cells were then washed twice with cold PBS, centrifuged at 3000 rpm for 5 minutes before we resuspended the cell pellet in 1 ml cold RIPA buffer (10 mM Tris pH 8.0, 1 mM EDTA, 140 mM NaCl, 1% Triton X-100, 0.1% SDS, 0.1% Na-Deoxycholate) supplemented with protease inhibitors and 125 mM glycine. Samples were snap-frozen in liquid nitrogen and stored at −80°C. Later, cells were thawed and incubated in a RIPA buffer (with 0.5% SDS) on ice for 15 min. The cells were vortexed before adding a RIPA buffer without SDS for a final concentration of 0.1% SDS in a volume of 300 μl. We sonicated the cells using a Bioruptor^®^ Standard with refrigerated sonication bath for 6-10 cycles of 30 sec ON and 30 sec OFF at high setting. Then the samples were briefly vortexed and debris was removed by centrifuging at top speed at 4°C for 15 min. Soluble chromatin supernatant equivalent to 6 μg per IP was pre-cleared using 20 μl of a 1:1 mix of protein A and protein G Dynabeads (Invitrogen). Overnight IP was performed using 1 μg of anti-H3K4me3 (Diagenode, C1541003-50), 2 μg anti-H3K27me3 (Diagenode, C15410195), and 2.5 μg anti-enhancer of zeste homolog 2 (EZH2) (Diagenode, C15200180). Antibody-protein complexes were immunoprecipitated with protein A/G Dynabeads for 3 hours at 4°C with rotation. Then the samples were washed for five minutes per wash with rotation; five times with 500 μl RIPA buffer supplemented with protease inhibitors, once with 500 μl LiCl wash buffer (250 mM LiCl, 10 mM Tris pH 8.0, 1 mM EDTA, 0.5% NP-40, 0.5% Na-Deoxycholate) supplemented with protease inhibitors, and once in 500 μl TE buffer (10 mM Tris-HCl pH 8.0, 1 mM EDTA). The beads were resuspended in 100 μl TE buffer and immunoprecipitated DNA was reverse cross-linked by adding 1 μl RNaseA [10 mg/ml] and incubated at 37°C for 30 min. Then we added 2.5 μl 20% SDS and 5 μl Proteinase K [10mg/ml] and incubated at 55°C for 1 hour, followed by purification using QIAquick PCR purification kit (Qiagen, 28104).

### Chromatin Immunoprecipitation qPCR (ChIP-qPCR)

For accurate fragment assessment, the shared chromatin was analyzed on a 1.5% agarose gel. Fragment size was optimized to be 200-500 bp. PCR primers were designed in the TSS region for the *SNHG26* neighbor genes; Translocase of Outer Mitochondrial Membrane 7 (*TOMM7)* and Family with Sequence Similarity 126 Member A (*FAM126A*) (Supplementary Table S5). As a control for successful IP, qPCR was performed using human positive and negative control qPCR primer sets from Active Motif (Supplementary Table S6). Furthermore, qPCR was performed on immunoprecipitated material and input chromatin. We added 2 μl ChIP DNA and 500 nM of each primer to SYBR Select Master Mix (Applied Biosystems) in three technical replicates. Target values from all qPCR samples were normalized with matched input DNA using the percent input method [100*2^ (Adjusted input - Ct (IP)]. The relative expression of *TOMM7* and *FAM126A* was normalized against *GAPDH*.

### Mitosox

We used MitoSOX™ Red Mitochondrial Superoxide Indicator (Thermo Fisher Scientific, M36008) to detect mitochondrial superoxide/reactive oxygen species (ROS formation). The MitoSOX red reagent permeates the cell membrane and selectively targets mitochondria where it is oxidized by superoxide. We seeded 40 000 cl/ml of HaCaT cells in a 24-well plate and transduced with gRNAs as previously described. After selection, the cells were washed twice with PBS and labeled with 5 μM Mitosox supplied in DMEM without phenol red (Thermo Fisher Scientific, 21063029) for 30 minutes at 37°C, protected from light. Then we washed the cells twice with pre-heated PBS and trypsinated for about 8 minutes before neutralizing with 500 μl DMEM without phenol red. The cells were centrifuged and re-suspended in 300 μl cold PBS, followed by BD FACS Canto flow cytometer (BD Bioscience) to detect the level of fluorescence with excitation/emission of 488/578. We used 50 μM Menadione as a positive control and no stained cells as a negative control.

### Total RNA sequencing (RNA-seq)

Total RNA was isolated using the *mir*Vana miRNA Isolation Kit (ThermoFisher Scientific, AM1560), according to the manufacturer’s protocol. RNA concentration was measured on a Qubit (Thermo fisher), whereas integrity and stability of the RNA samples were assessed by using an Agilent 2100 Bioanalyzer (Agilent Technologies). The ribosomal RNA (rRNA) was removed using RiboCop rRNA Depletion Kit for Human/Mouse/Rat V2 (Lexogen), followed by preparation of RNA-seq libraries using the CORALL total RNA library prep (Lexogen), according to the manufacturer’s instructions. The samples were sequenced on an Illumina NextSeq HighOutput v2.5 flow cell using the Illumina NextSeq 500 Sequencing System.

### RNA-seq data analysis

FASTQ files were filtered and trimmed (fastp v0.20.0) and transcript counts were generated using quasi alignment (Salmon v1.3.0) to the transcriptome reference sequence (Ensembl, GRCh38 release 92). We imported the transcript sequences into the R statistical software and aggregated them to gene counts using the tximport (v1.14.0) bioconductor package. To compare gene expression between samples, the genes with expression of less than 1 count per million in at least 50% of the samples were filtered out. The count matrices were transformed using bioconductor package limma (Ritchie et al., 2015) combined with voom transformation (Law et al., 2014). In the limma framework, gene expression was modelled as a function of four binary variables representing cell line (CRISPRi/a) combined with treatment (*SNH26* or control gRNA). Three contrasts were analyzed to identify differentially expressed genes: (i) differences between gRNA treatment in CRISPRi, (ii) in CRISPRa, and (iii) and interactions between gRNA treatments in CRISPRi and CRISPRa (i.e. differences between difference (i) and (ii)). Differentially expressed genes were identified using the topTable function of limma and Benjamini-Hochberg method was used to correct for multiple testing, and genes with *p*-values <0.05 were considered significantly differentially expressed.

## Acknowledgement

This work was supported by the Norwegian Cancer Society (grant number 2278701) and the Research Council of Norway (grant number 230338). The library preparation and sequencing were provided by the Genomics Core Facility (GCF), Norwegian University of Science and Technology (NTNU). GCF is funded by the Faculty of Medicine and Health Sciences at NTNU and Central Norway Regional Health Authority. The light microscopy was provided by the Cellular and Molecular Imaging Core Facility (CMIC), NTNU. CMIC is funded by the Faculty of Medicine at NTNU and Central Norway Regional Health Authority.

## Author contribution

HS designed and performed all imaging and wet lab experiments, contributed with statistical analysis, and wrote the manuscript. SAH contributed with statistical analysis, generated the figures, and was a major contributor in editing the manuscript. KC analyzed the RNA-seq data. NBL performed all FACS analysis. PAA contributed with the planning and design of CRISPR experiments. BS contributed to the planning of RNA FISH experiments and performed the image analysis. PS contributed to the study design, supervised the project, and edited the manuscript. All authors read and approved the final manuscript.

## Data availability

The RNA-seq dataset generated and analyzed during the current study is available in the European Nucleotide Archive (https://www.ebi.ac.uk/ena/) under the accession number of: PRJEB43155

## Supplementary Data

**Supplementary file 1:** Supplementary Figures and Tables

**Supplementary dataset 1**: A list of 1623 genes that were differentially expressed (*p-*value < 0.05) in CRISPRa vs. CRISPRi in HaCaT cells.

## Supplementary Figures

**Supplementary Figure S1:**
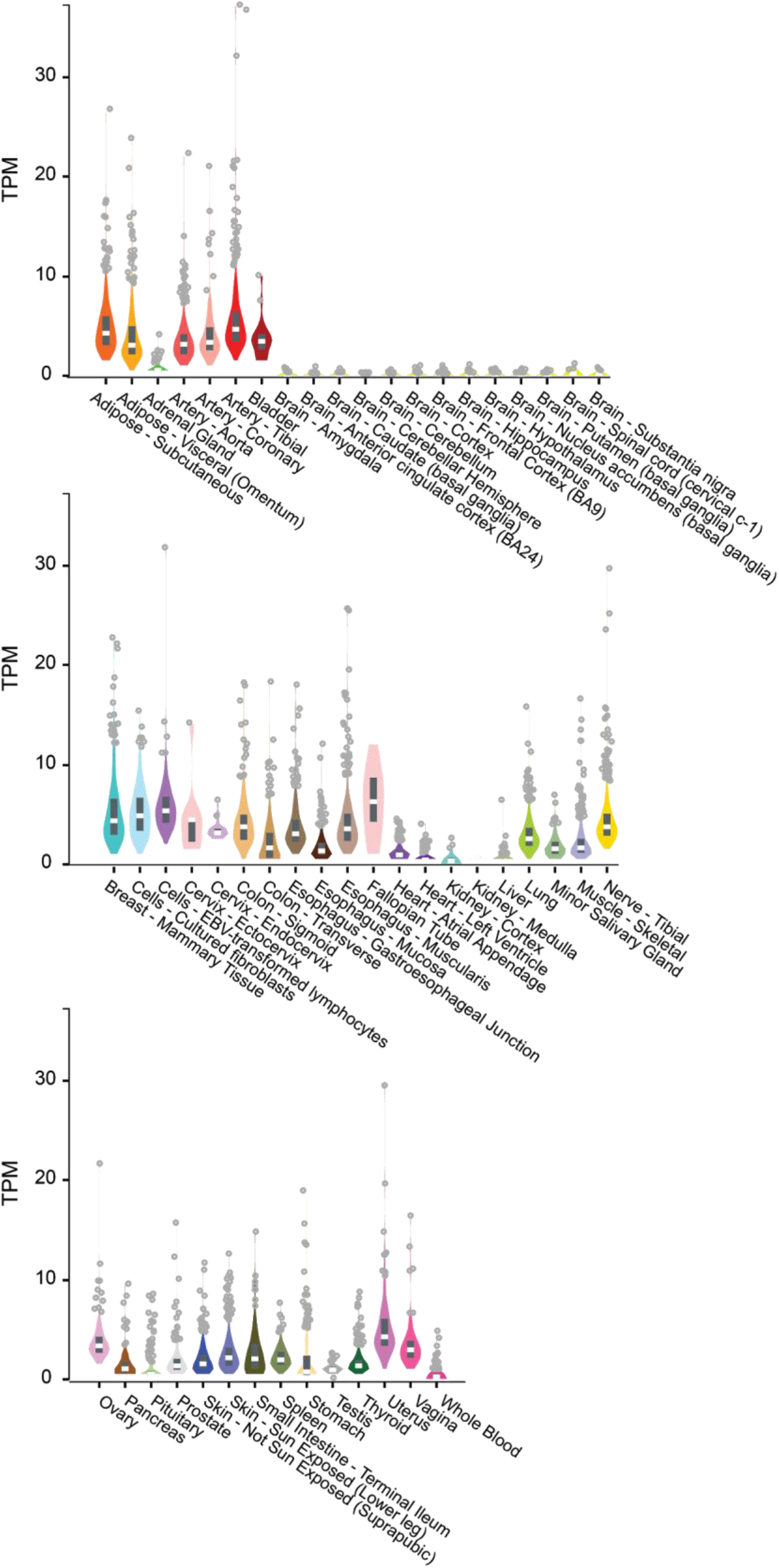
Tissue expression (transcript per kilobase million, TPM) for *SNHG26* (ENSG00000228649) from the Genotype-Tissue Expression (GTEx) project.

**Supplementary Figure S2:**
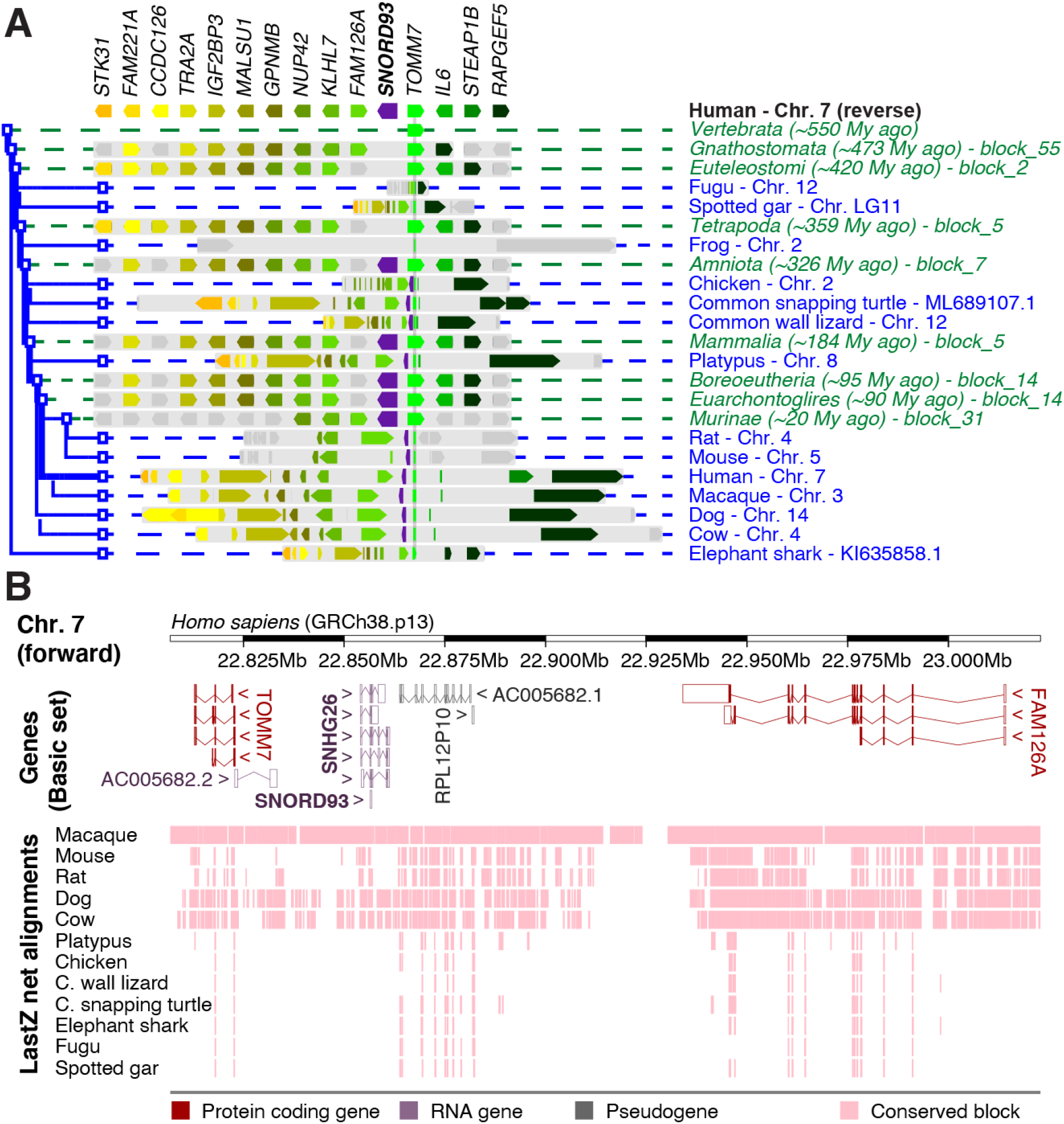
The *TOMM7-SNORD93-FAM126A* locus is conserved in Amniota. (**A**) Evolutionary conservation of genes in the locus. The view is centered on *TOMM7*, which is conserved in Vertebrata, and shows conservation of 14 neighboring human genes (color coded) in syntenic blocks (light gray) from selected species (blue) and ancestors (green). Genes in species are shown on a comparable genomic scale; genes in ancestors are shown as schematics. Other genes than those that are homologous to the 15 human genes from the locus are shown in grey. Data from Genomicus [1], modified to include *SNORD93* conservation. (**B**) Overview of the human *TOMM7-SNORD93-FAM126A* locus. ”Genes’’ show transcripts from the GENCODE Basic set (Ensembl release 102); “LastZ net alignments” show conserved sequence blocks in the species from (A).

**Supplementary Figure S3:**
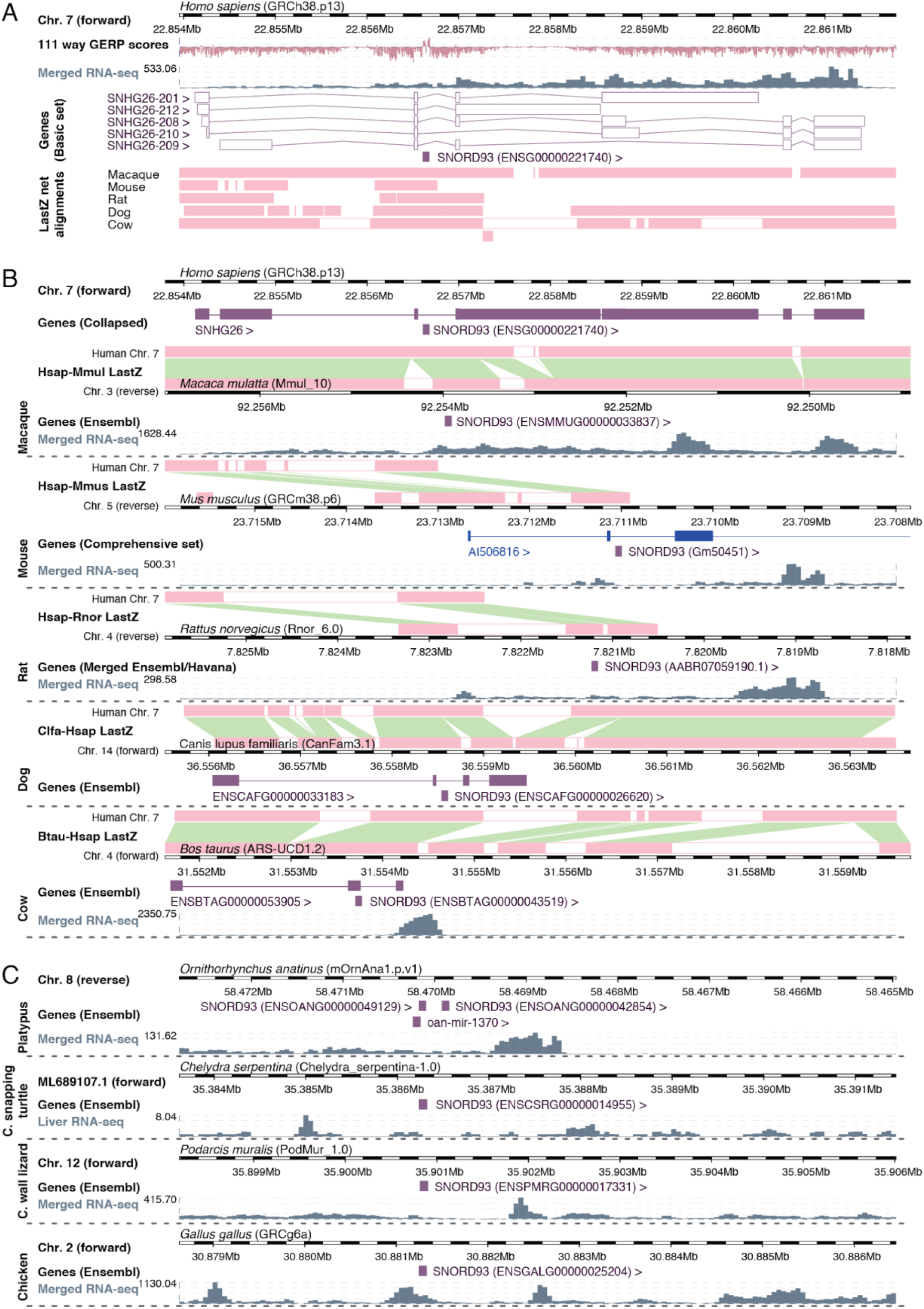
Comparative genomics and RNA-seq data indicate conserved sequence and expression for *SNHG26*. (**A**) Overview of the human *SNHG26* locus. The genome browser tracks show Genomic Evolutionary Rate Profiling (GERP) scores, RNA-seq read coverage from the Human BodyMap 2.0 project (ArrayExpress ID: E-MTAB-513), transcripts from the GENCODE Basic set (Ensembl release 102), and sequence blocks that are conserved in the species from Figure S2A. The genomic blocks containing *SNHG26* exons 1-3 are largely conserved in all five species, whereas the last exons appear lost in rodents. (**B**) Sequence block conservation (pairwise LastZ maps), annotated genes (Genes), and RNA-seq read coverage (RNA-seq) within the *SNHG26* homologous regions for the species in (A). Mouse, dog, and cow each have an annotated non-coding RNA (ncRNA) mapping to the *SNHG26* homologous regions. Each species’s RNA-seq data show transcription within the locus. (**C**) Annotated genes (Genes) and RNA-seq read coverage (RNA-seq) in the genomic regions surrounding the *SNORD93* homologues in platypus, chicken, turtle, and lizard. Each species’s RNA-seq data show transcription within the overall locus.

**Supplementary Figure S4:**
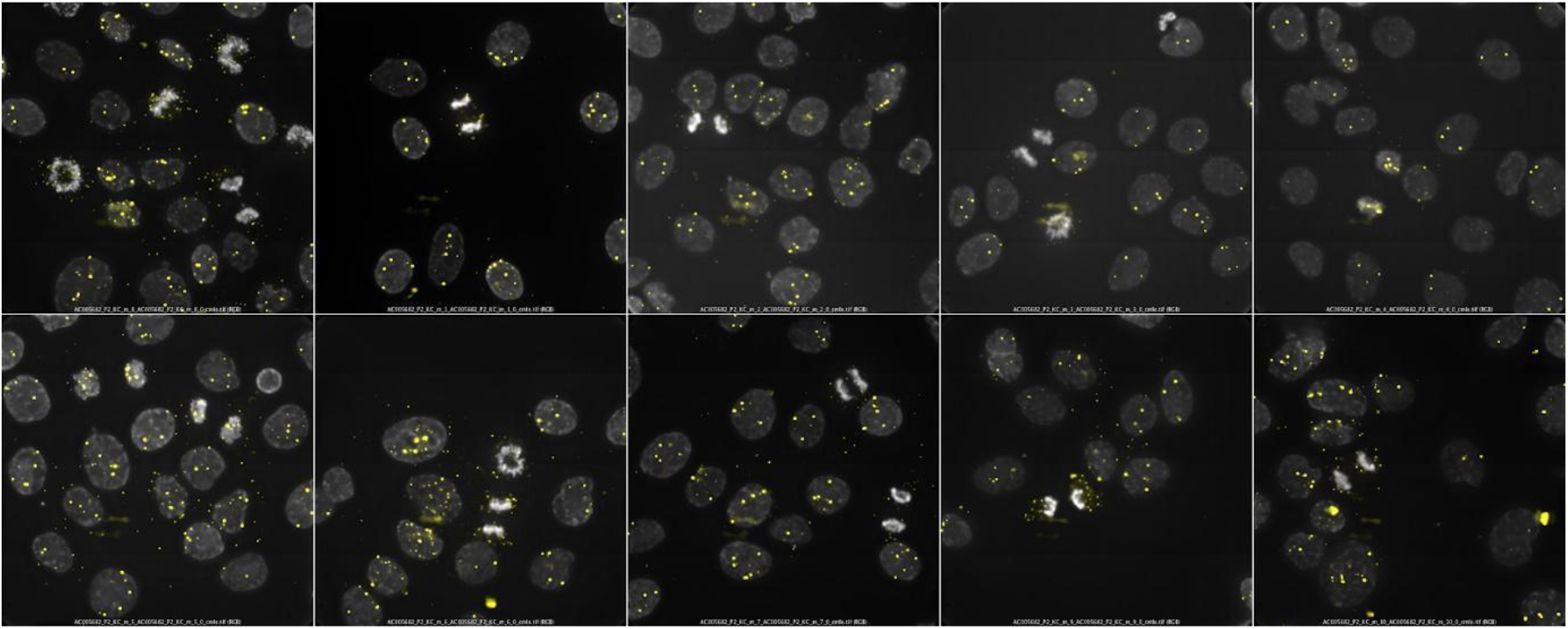
Mitotic HaCaT cells. The localization of *SNHG26* (yellow) in mitotic fixed HaCaT cells stained with DAPI (grey).

**Supplementary Figure S5:**
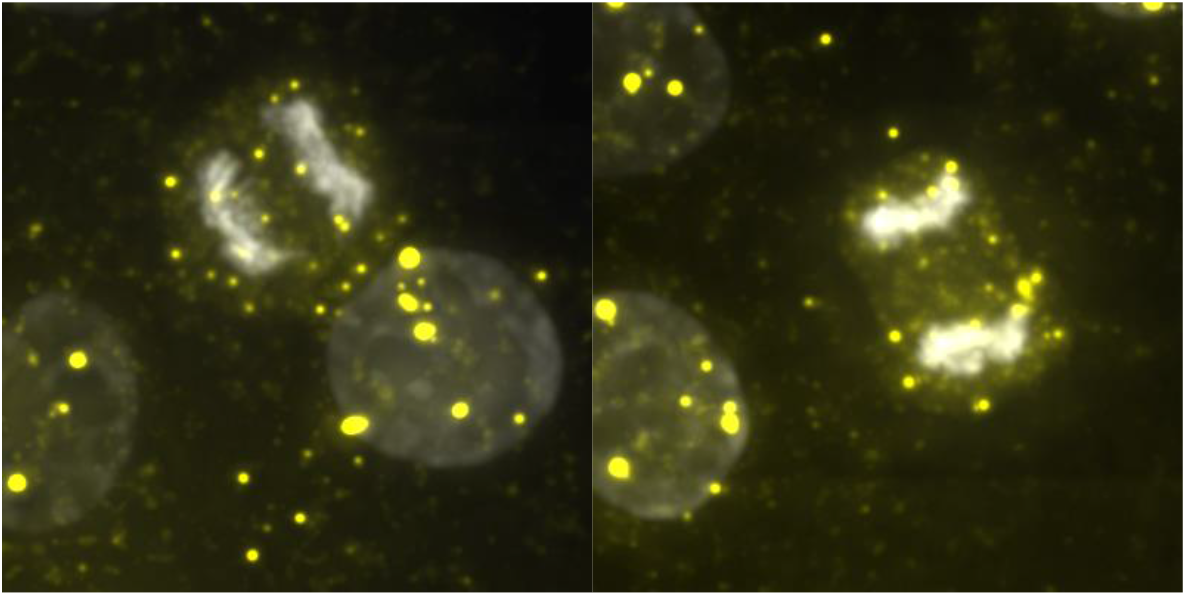
Mitotic HaCaT cells with low-intensity spots. The localization of *SNHG26* (yellow) in mitotic fixed HaCaT cells stained with DAPI (grey) with spots between 100-1500 intensities.

**Supplementary Figure S6:**
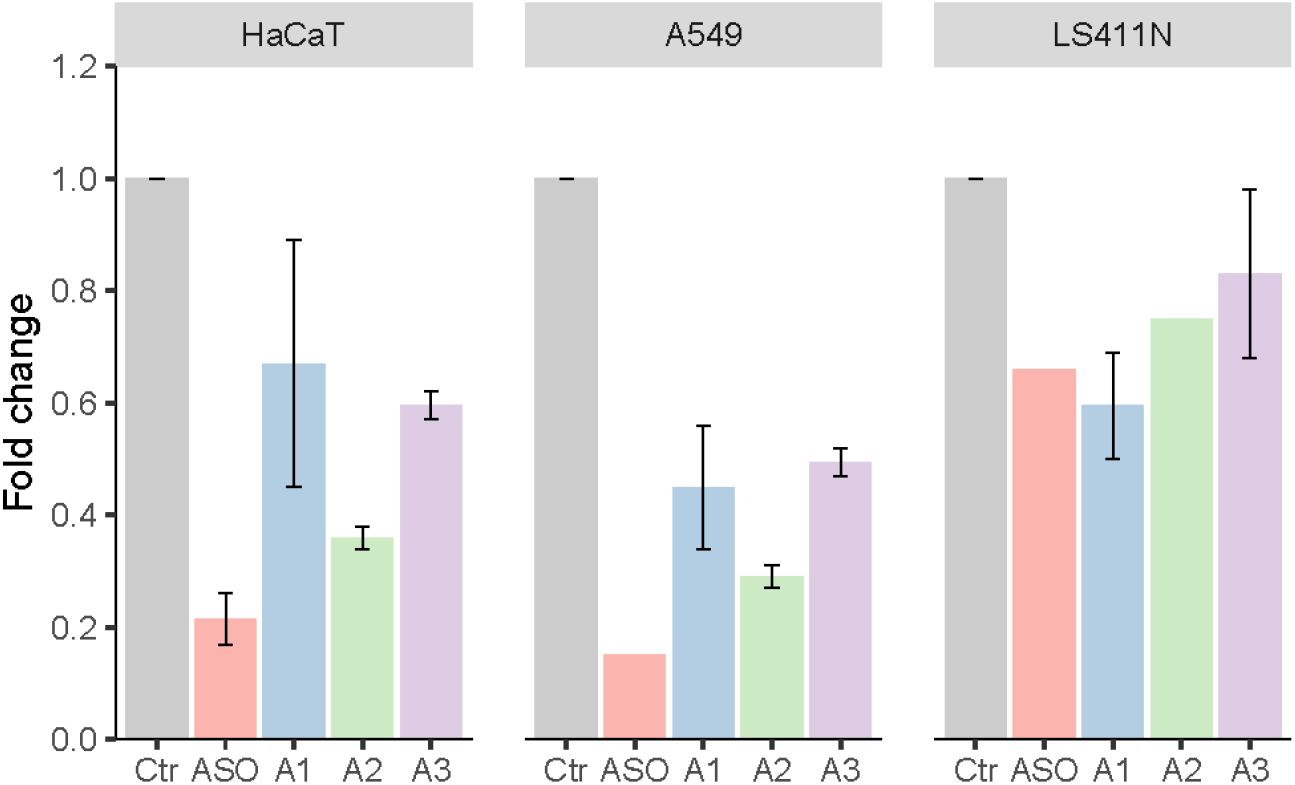
Percentage downregulation of *SNHG26* by using ASO and three siRNAs with different target sequences. Data are presented as fold change expressions of *SNHG26* following ASO or siRNA treatment relative to control-treated cells as measured by RT-qPCR in HaCaT, A549, and LS411N cells. Bars and error bars represent mean and standard error of mean (SEM) for HaCaT (n=2), A549 (n=1 for ASO and n=2 for siRNAs A1-A3), LS411N (n=1 for ASO and A1, n=2 for A2 and A3).

**Supplementary Figure S7:**
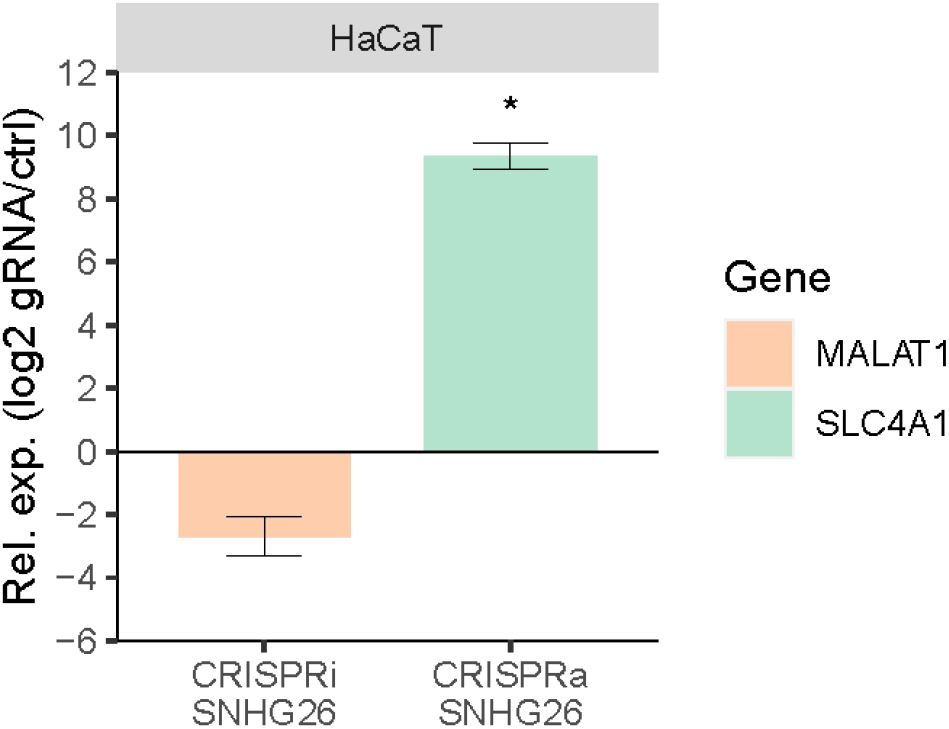
The relative expression level of *MALAT1* and *SLC4A1* as measured by RT-qPCR in response to CRISPRi/a of *SNHG26* **in HaCaT cells.** Data are presented as fold change expressions of cells transduced with target-specific gRNA relative to control gRNA. Bars and error bars are mean and SEM of two independent replicates. Significant differences were determined by Student’s *t*-test (unpaired, two-tailed), assuming equal variances (\**p* ≤ 0.05).

**Supplementary Figure S8:**
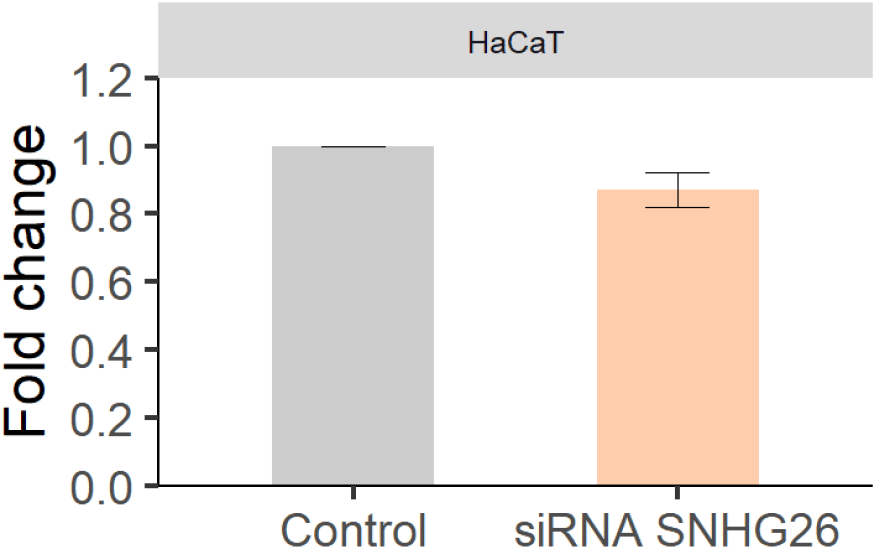
Effects of siRNA-mediated knockdown of *SNHG26* on metabolic activity in HaCaT cells, as measured by XTT. Data are presented as normalized absorbance (A_465 nm_ – A_630 nm_) for cells transfected with *SNHG26*-specific siRNA and control siRNA. Bars and error bars are mean and SEM of three independent replicates. *P*-value was calculated by Student’s t-test, (unpaired, two-tailed) assuming equal variances (*p* = 0.13).

**Supplementary Figure S9:**
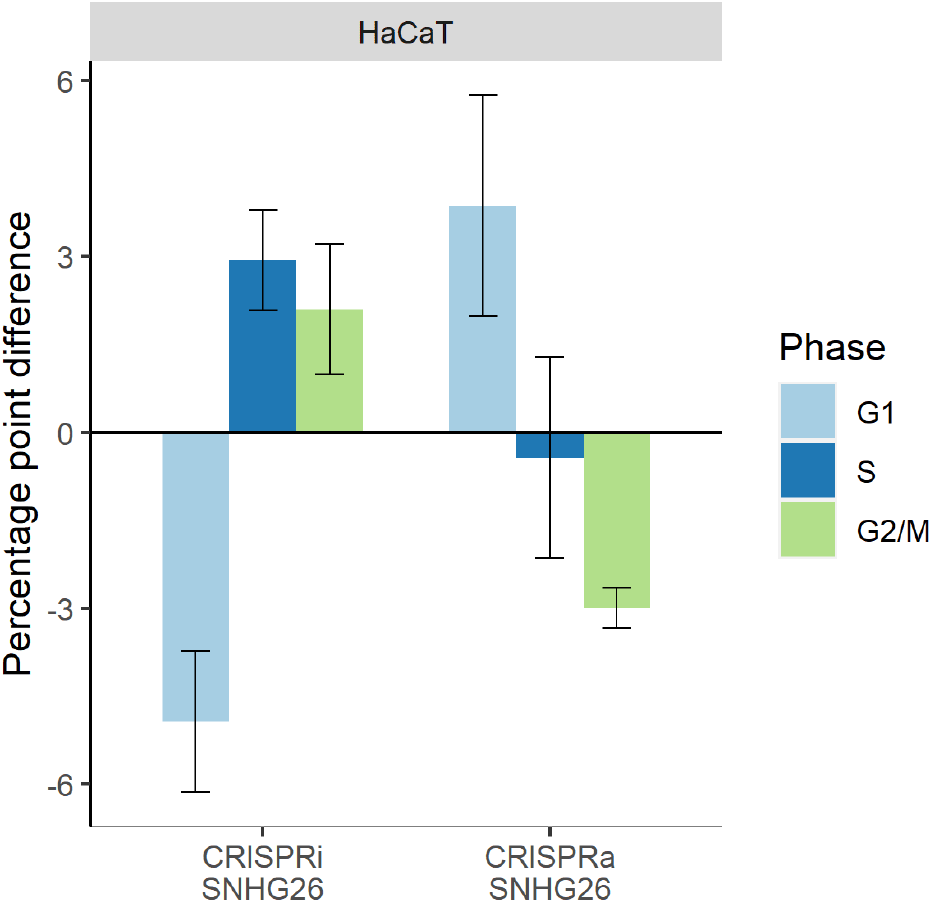
The distribution of cells in G1, S, and G2/M cell cycle phases in response to CRISPRi/a modulation of *SNHG26* in HaCaT cells harvested and re-seeded followed by cell cycle assay eight days after gRNA transduction. Data are the difference in percentages of G1, S, and G2/M cells between cells transduced with target-specific gRNA to those transduced with a control gRNA. Bars and error bars are mean and SEM of three independent replicates. ANOVA *p*-values were calculated from a hierarchical, linear model (G1: 2.32e-4, S: 0.082, and G2/M: 0.0131).

**Supplementary Figure S10:**
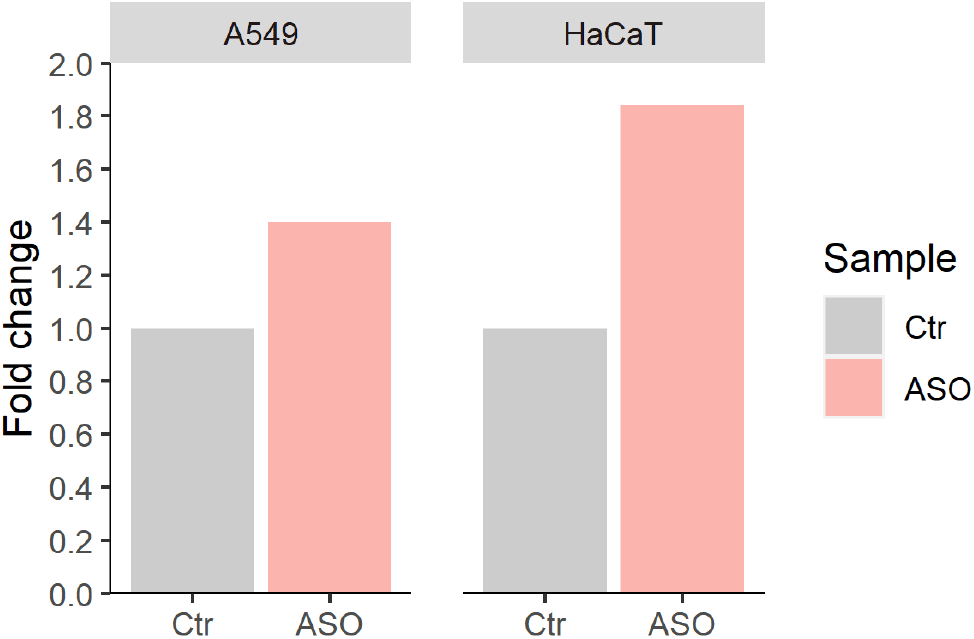
We used Mitosox and FACS to determine whether ASO-mediated modulation of *SNHG26* affected mitochondrial stress in HaCaT and A549 cells (n=1). Data are presented as fold change fluorescence intensity of cells transduced with target-specific gRNA relative to control gRNA.

**Supplementary Figure S11:**
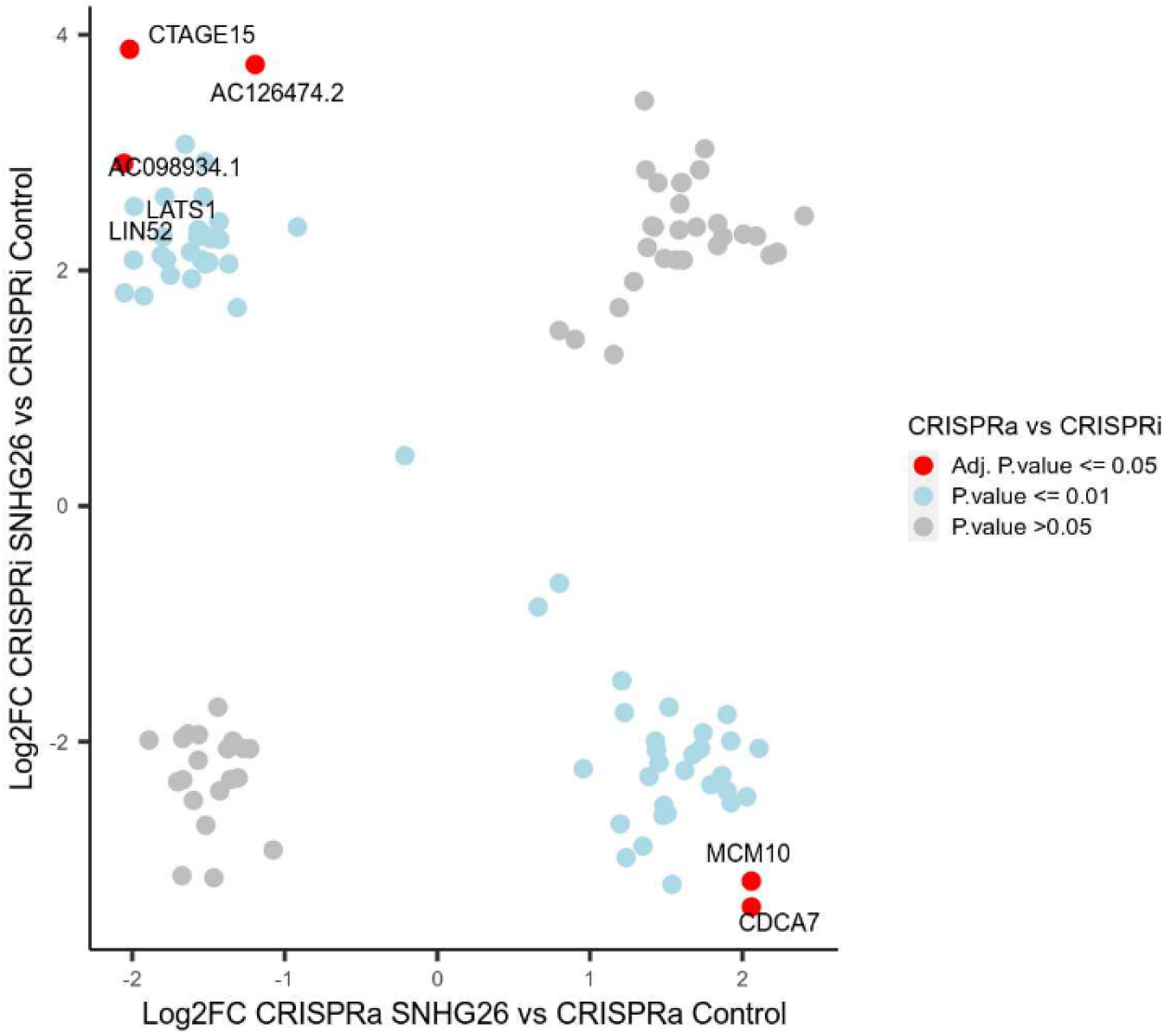
A total of 110 differentially expressed genes with *P*-value < 0.05 in both CRISPRi vs. Control and CRISPRa vs. Control. Legend shows their adjusted *P*-value in the CRISPRa vs. CRISPRi comparison. The five significant genes marked in red were differentially expressed in the opposite direction in CRISPRa compared to CRISPRi (adjusted *P*-value < 0.05).

## Supplementary Tables

**Supplementary Table S1:**
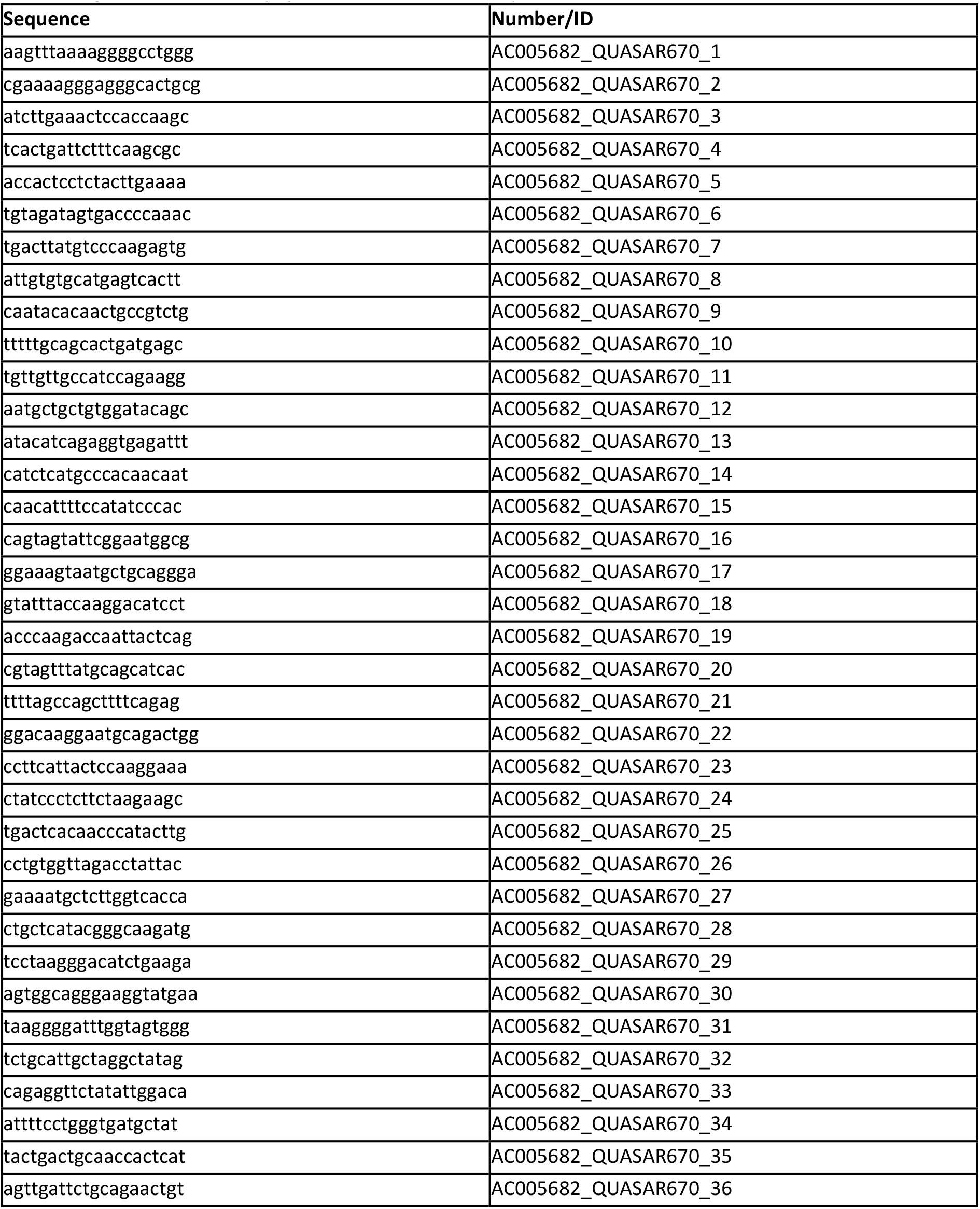
Custom Stellaris FISH probes. The probes were ordered from Biosearch Technologies and were conjugated to a Quasar670 dye in the 3′end.

**Supplementary Table S2:**
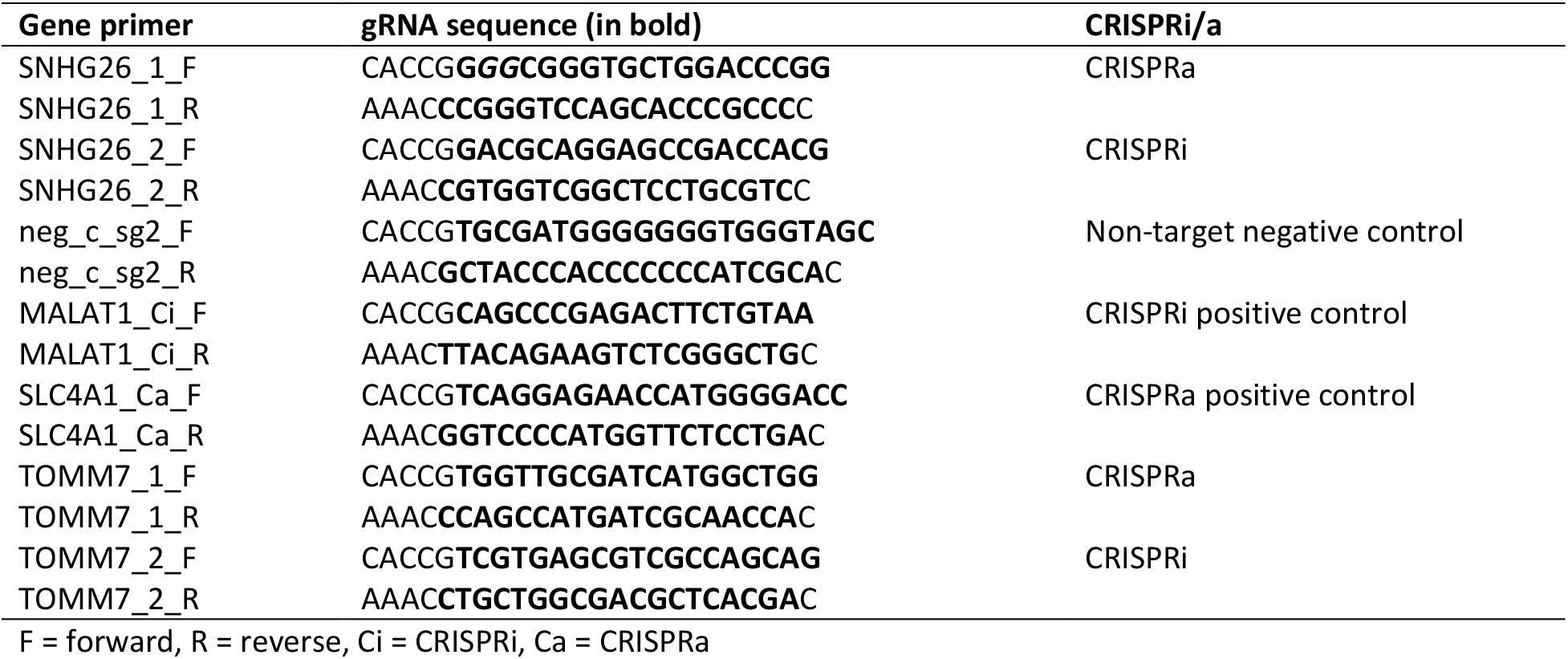
Oligos used for guide RNA (gRNA) cloning.

**Supplementary Table S3:**
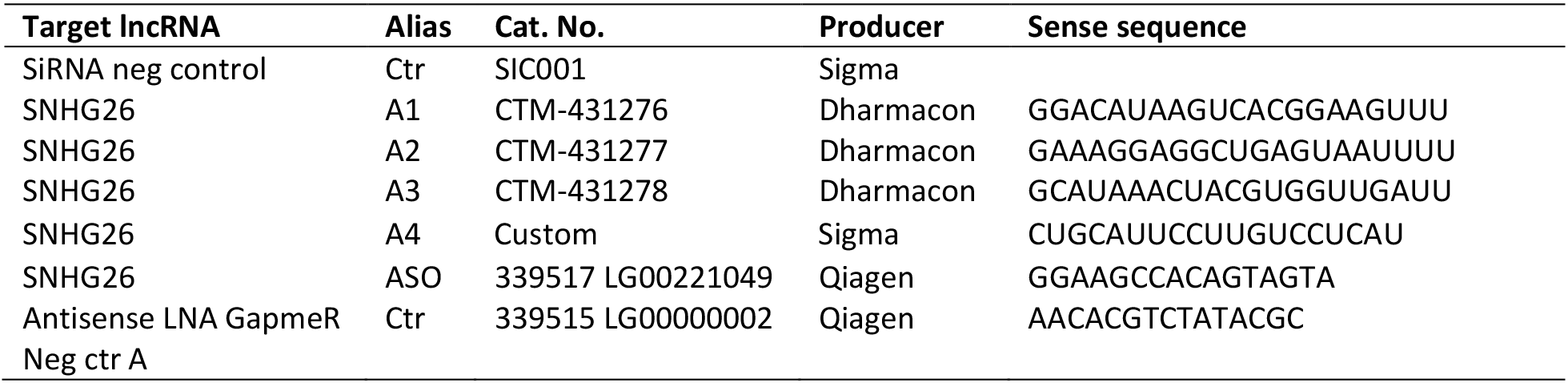
Antisense oligo and siRNAs used in RNA interference experiments.

**Supplementary Table S4:**
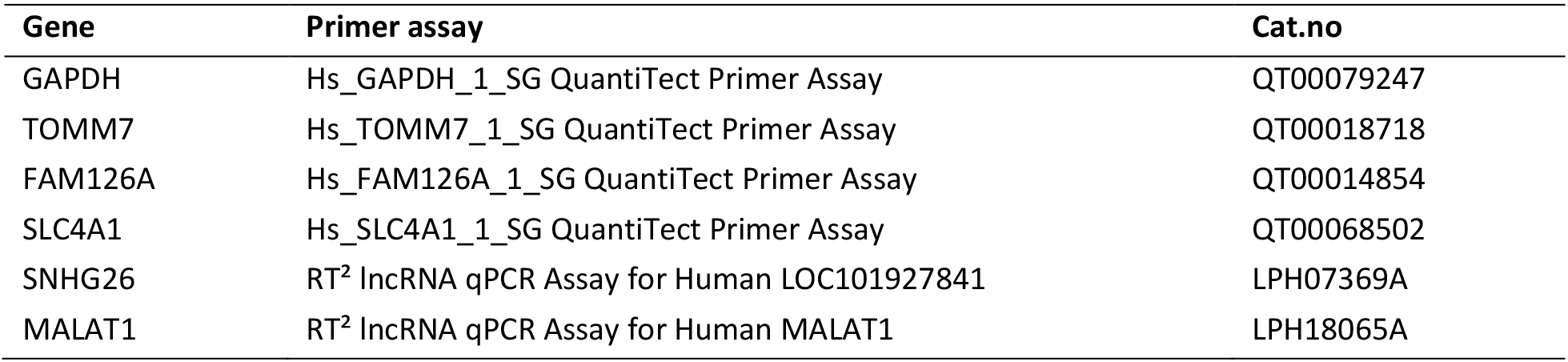
Primers used for RT-qPCR. Qiagen QuantiTect Primer Assays (249900) were used for mRNA expression analysis and Qiagen RT^2^ lncRNA. PCR assays (330701) were used for lncRNA expression analysis

| Gene | Primer assay | Cat.no |
| --- | --- | --- |
| GAPDH | Hs_GAPDH_1_SG QuantiTect Primer Assay | QT00079247 |
| TOMM7 | Hs_TOMM7_1_SG QuantiTect Primer Assay | QT00018718 |
| FAM126A | Hs_FAM126A_1_SG QuantiTect Primer Assay | QT00014854 |
| SLC4A1 | Hs_SLC4A1_1_SG QuantiTect Primer Assay | QT00068502 |
| SNHG26 | RT<sup>2</sup> lncRNA qPCR Assay for Human LOC101927841 | LPH07369A |
| MALAT1 | RT<sup>2</sup> lncRNA qPCR Assay for Human MALAT1 | LPH18065A |

**Supplementary Table S5:**
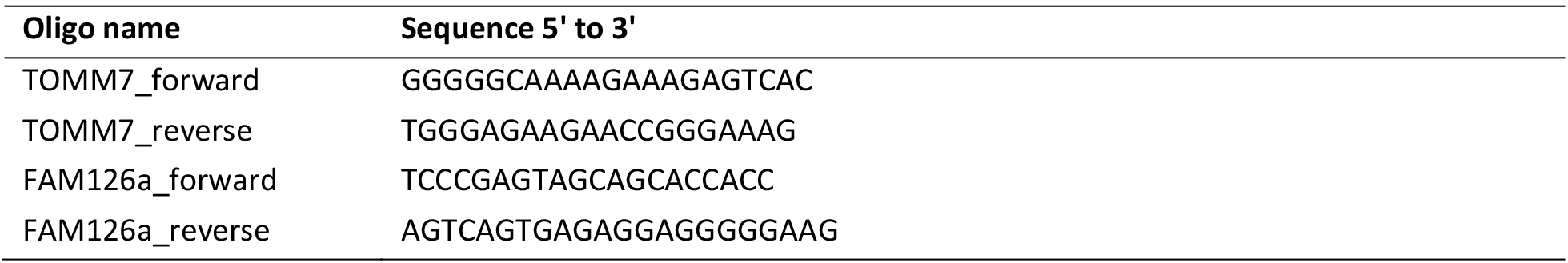
Custom primers targeting transcriptional start site (TSS) used for RT-qPCR. Custom DNA oligos were purchased from Sigma-Aldrich.

**Supplementary Table S6:**
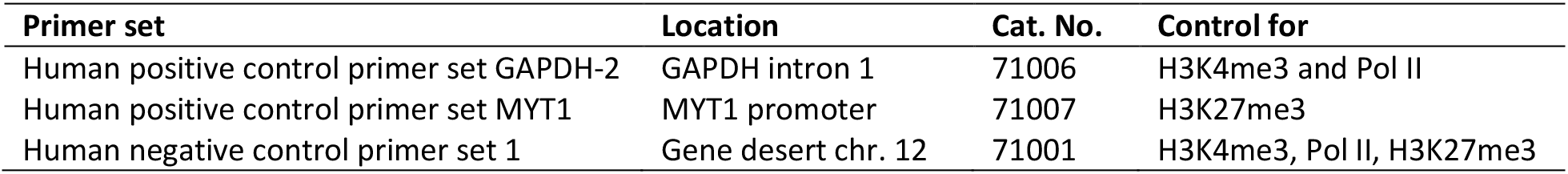
Human control qPCR primer sets from Active Motif. Human Control qPCR Primer Sets are designed to serve as positive or negative ChIP controls when performing chromatin immunoprecipitation (ChIP) with human samples. Each primer set is a mixture of forward and reverse primers that have been validated for qPCR and endpoint PCR of ChIP samples from multiple human cell lines.

## Supplementary Dataset

**Supplementary Dataset S1: A list of 1623 genes that were differentially expressed (*p-*value < 0.05) in CRISPRa vs. CRISPRi.**The results of Limma toptable function from individual (CRISPRa vs. Control and CRISPRi vs. Control) and joint analysis (CRISPRa vs. CRISPRi).

